# Gfi1 controls the formation of exhausted effector-like CD8 T cells during chronic infection and cancer

**DOI:** 10.1101/2024.04.18.579535

**Authors:** Oluwagbemiga A Ojo, Hongxing Shen, Jennifer T Ingram, James A Bonner, Robert S Welner, Georges Lacaud, Allan J Zajac, Lewis Z Shi

## Abstract

During chronic infections and tumor progression, CD8 T cells gradually lose their effector functions and become exhausted. These exhausted CD8 T cells are heterogeneous and comprised of different subsets, including self-renewing progenitors that give rise to Ly108^−^ CX_3_CR1^+^ effector-like cells. Generation of these effector-like cells is essential for the control of chronic infections and tumors, albeit limited. However, the precise cues and mechanisms directing the formation and maintenance of exhausted effector-like are incompletely understood. Using genetic mouse models challenged with LCMV Clone 13 or syngeneic tumors, we show that the expression of a transcriptional repressor, growth factor independent 1 (Gfi1) is dynamically regulated in exhausted CD8 T cells, which in turn regulates the formation of exhausted effector-like cells. Gfi1 deletion in T cells dysregulates the chromatin accessibility and transcriptomic programs associated with the differentiation of LCMV Clone 13-specific CD8 T cell exhaustion, preventing the formation of effector-like and terminally exhausted cells while maintaining progenitors and a newly identified Ly108^+^CX_3_CR1^+^ state. These Ly108^+^CX_3_CR1^+^ cells have a distinct chromatin profile and may represent an alternative target for therapeutic interventions to combat chronic infections and cancer. In sum, we show that Gfi1 is a critical regulator of the formation of exhausted effector-like cells.

## Introduction

During chronic infections such as hepatitis B virus in humans and lymphocytic choriomeningitis virus CL-13 (hereafter, LCMV CL-13) in mice^(*1*)^ and tumor progression, persistent antigen stimulation causes CD8 T cells to progressively lose their effector functions, in a process termed T cell exhaustion^(*2–6*)^. Therefore, advancing our understanding of T cell exhaustion is important and would improve immunotherapeutic interventions for chronic infections and cancer.

Exhausted CD8 T cells are heterogeneous and maintained by self-renewing TCF1^+^Ly108^+^ progenitor cells^(*4, 7–11*)^. Immune checkpoint blockade (ICB) therapies (e.g., anti-CTLA-4 and anti-PD-1) act on these progenitors, induce their proliferation, and mediate therapeutic effects^(*4, 7, 8, 12*)^. Proliferating progenitors adopt either a Ly108^−^CX_3_CR1^+^ effector-like or a Ly108^−^CX_3_CR1^−^ terminally exhausted state^(*2, 4, 7, 8, 11–16*)^, with the former possessing greater cytolytic functions and exerting control of cancer and chronic viral infections^(*16, 17*)^ to certain extent. Accumulating evidence indicates that generation of these various exhausted subsets requires substantial chromatin remodeling in progenitor cells^(*10, 18–21*)^, a process orchestrated by a complex network of transcription factors (TFs) and signaling pathways. While TOX is critical for the overall development and maintenance of exhausted CD8 T cells^(*22–25*)^, other TFs are also involved. TCF1 is necessary for the formation and maintenance of progenitors, and this is further reinforced by Bach2 and PBAF^(*9, 10, 19–21, 26*)^. Batf and Zeb2 facilitate the formation of the effector-like and terminally exhausted subsets, but deletion of Batf and Zeb2 does not prevent their formation^(*9, 13, 14, 18*)^, suggesting that other unidentified TFs/regulators also play an important role in this process.

Growth factor-independent 1 (Gfi1) is a transcriptional repressor, consisting of a SNAG domain^(*27, 28*)^ and six zinc-finger motifs of which 3-5 are crucial for its DNA binding activity^(*28*)^. Gfi1 represses target genes^(*29–31*)^ through recruitment of epigenetic enzymes such as histone deacetylases (HDACs) and demethylases (KDMs)^(*32, 33*)^ and by participating in several chromatin modifying complexes such as CoREST, NuRD, SWI/SNF, and CtBP^(*34*)^. We and others show that Gfi1 orchestrates the development of thymocytes, and its absence favors the formation of single positive CD8 T thymocytes^(*35–40*)^, partially dependent on its suppression of Foxo1, Klf2, Irf4, *etc.*^(*36, 39*)^. Given this, the role of Gfi1 in peripheral mature CD8 T cells is unknown, likely due to low Gfi1 expression in naïve CD8 T cells, although it is greatly increased upon antigen stimulation^(*41, 42*)^.

Considering that Gfi1 intricately interacts with other TFs, epigenetic enzymes, and chromatin modifying complexes, here we investigate whether and how Gfi1 modulates CD8 T cell exhaustion during chronic infection and cancer. Using Gfi1-tdTomato reporter mice infected with LCMV CL-13, we found that Gfi1 expression was dynamically regulated in the exhausted antigen-specific CD8 T cells: first emerged in TCF1^+^ progenitors, downregulated in Ly108^+^CX_3_CR1^+^ cells, and expressed in Ly108^−^CX_3_CR1^+^ effector-like and Ly108^−^CX_3_CR1^−^ terminally exhausted cells. Specific deletion of Gfi1 in T cells (Gfi1^cKO^) blocked the formation of effector-like and terminally exhausted CD8^+^ T cells, with accompanying accumulation of progenitors and Ly108^+^CX_3_CR1^+^ cells. Using the assay for transposase-accessible chromatin with high-throughput sequencing (ATAC-seq) and RNA-seq to assess genome-wide chromatin accessibility and transcriptomes, we revealed severely dysregulated chromatin structures and transcriptomic programs in Gfi1^cKO^ progenitors and Ly108^+^CX_3_CR1^+^ cells. Additional studies with *in vivo* deletion of CD4 T cells and chimeric mice indicated that the arrested development of exhausted Gfi1^cKO^ CD8 T cell at the Ly108^+^CX_3_CR1^+^ stage was a cell-autonomous effect. Similarly, tumor infiltrating Gfi1^cKO^ CD8 T cells (CD8 TILs) were impaired to form terminally differentiated effector cells from progenitors, and Gfi1^cKO^ mice bearing tumor were not responsive to ICB anti-CTLA-4, coupled with reduced effector function (IFN-γ production) of CD8 TILs. Taken together, we show that Gfi1, by sustaining appropriate chromatin accessibility and transcriptomes, controls the generation of effector and terminally exhausted cells from progenitors during chronic infection and tumor progression.

## Results

### Gfi1 expression in CD8 T cells is dynamically regulated during chronic infection

To monitor Gfi1 expression in CD8 T cells in response to chronic viral infection, we employed the Gfi1 reporter mouse model, wherein endogenous Gfi1 expression is measured by tdTomato^(*43*)^ (hereafter, Gfi1^tdTomato^). Gfi1^tdTomato^ mice were infected with LCMV CL-13, as we previously described^(*2*)^, and on days 5, 8, 15, and 30 post-infection, splenocytes were stained with LCMV-D^b^-GP33 tetramer to detect LCMV GP33 antigen-specific CD8 T cells. While Gfi1 was not expressed in LCMV GP33^+^ CD8 T cells during the earliest phase of the chronic infection (days 5 and 8), its expression gradually increased at later times and reached the peak level on day 30 (Fig. 1A). We then asked if Gfi1 was preferentially expressed on specific exhausted subsets as they emerged during chronic viral infection. As shown in Fig. S1A, we did not see substantial Gfi1 expression in any of the subsets, defined by Ly108 (surrogate marker for TCF1^(*6*)^) and CX_3_CR1 (an effector cell marker^(*44*)^) between days 5-8 following infection. As the chronic phase kicked in around day 15, Gfi1 expression was increased on Ly108^+^CX_3_CR1^−^ progenitors but largely unaltered on the effector-like (Ly108^−^CX_3_CR1^+^) and terminally exhausted cells (Ly108^−^CX_3_CR1). Of note, the progenitor cells were present from day 5-8, but they did not express Gfi1, suggesting that the development of exhaustion induced Gfi1 expression.

**Figure 1.**
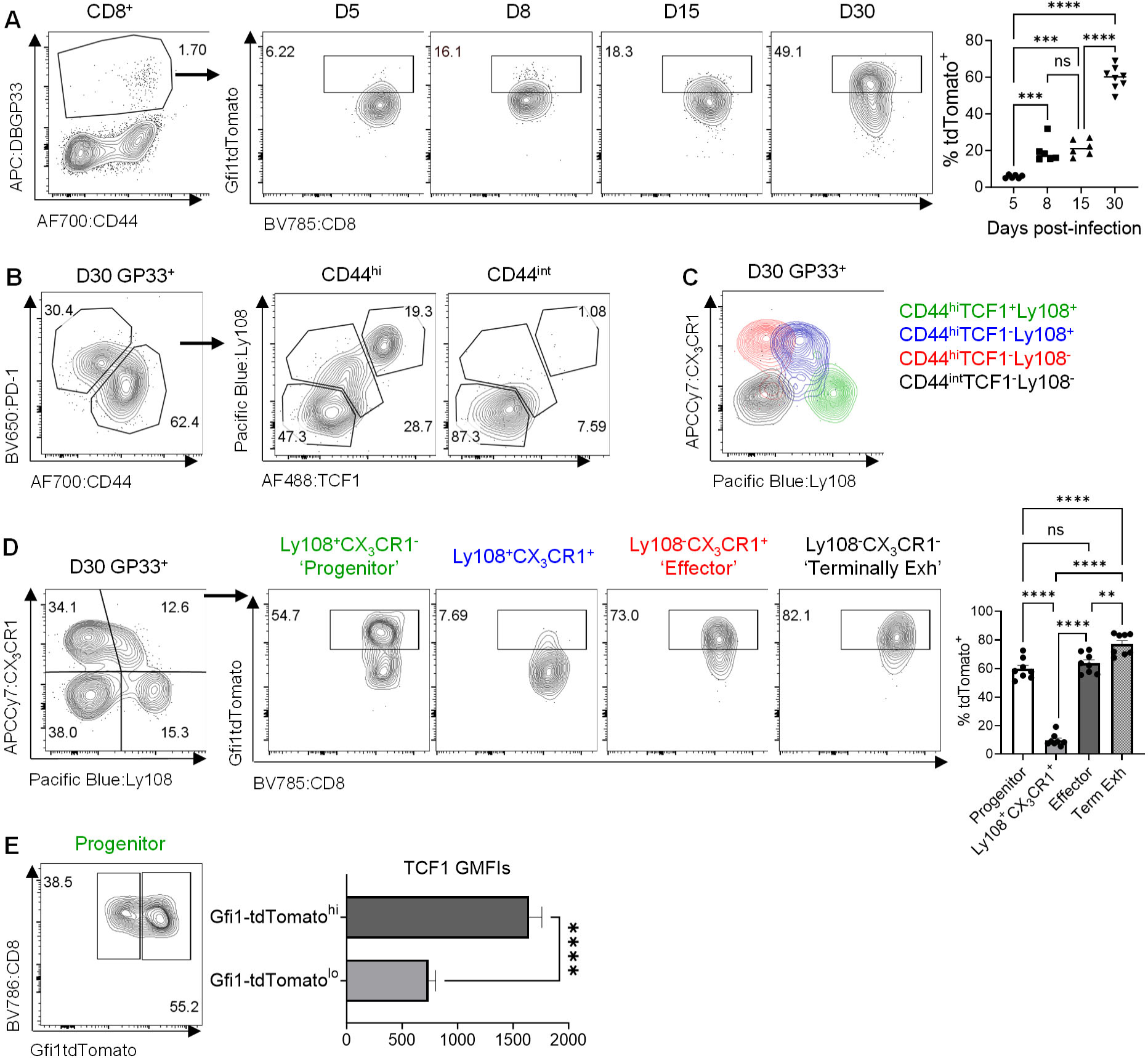
Gfi1 is differentially expressed in exhausted CD8 subsets during chronic infection. **A.** Splenocytes from Gfi1^tdTomato^ reporter mice infected with LCMV Cl-13 were stained for GP33 tetramer and analyzed for tdTomato (Gfi1) expression days 5, 8, 15 and 30 post-infection. **B.** Identification of the fourth distinct population (CD44^hi^TCF1^−^Ly108^+^) of exhausted CD8^+^ T cells on day 30 based on CD44, PD1, TCF1 and Ly108 expression, in addition to the CD44^hi^TCF1^+^Ly108^+^, CD44^hi^TCF1^−^Ly108^−^, and CD44^int^TCF1^−^Ly108^−^ subsets. **C.** A clear demarcation of the four subsets from **B** with Ly108 and CX_3_CR1 on day 30 post-infection. **D.** Gfi1 expression on defined CD8^+^ T cell subsets in **C**. **E.** GMFIs of TCF1 on tdTomato^hi^ versus tdTomato^lo^ sub-populations in the progenitor subset from **D**. Data were from 2 independent experiments with n=7-8, depicted as mean ± s.e.m. ns, P≥0.05; **, P<0.01; ***, P<0.001; ****, P<0.0001. Data in **A** and **D** were analyzed by one-way ANOVA with Tukey’s or Sidak’s post-hoc test; data in **E** were analyzed with paired two-tailed Student’s t test.

The exhausted T cell pool is established by 30 days post-infection and comprised of 3 well-defined subsets^(*45*)^. These subsets can be identified by PD-1, CD44, Ly108 and TCF1 (Fig. 1B). Specifically, CD44 expression diminished as CD8 T cells became terminally exhausted^(*2*)^, dividing GP33^+^ LCMV-specific cells to CD44^hi^ and CD44^int^ cells. While CD44^int^ were homogeneously Ly108^−^TCF1^−^ terminally exhausted cells, the CD44^hi^ cells contained 3 sub-populations: Ly108^+^TCF1^+^, Ly108^+^TCF1^−^ and Ly108^−^TCF1^−^ cells (Fig. 1B). We then used CX_3_CR1 to identify the effector-like subsets of exhausted CD8^+^ T cells (Fig. 1C). As expected, both CD44^int^Ly108^−^ TCF1^−^ terminally exhausted and CD44^hi^Ly108^+^TCF1^+^ progenitor cells expressed CX_3_CR1 at low levels, whereas CD44^hi^Ly108TCF1^−^ effector-like cells expressed it highly. Intriguingly, our gating (Fig. 1B&C) and back-gating (Fig. S1B) strategies also identified a fourth distinct population, the CD44^hi^Ly108^+^TCF1^−^ cells that expressed CX_3_CR1 and Ly108 at intermediate levels, which have not been previously well-described (Fig. S1C). We analyzed Gfi1 expression on these 4 subsets. As shown in Fig. 1D, Gfi1 was highly expressed in the Ly108^+^CX_3_CR1^−^ progenitors, Ly108^−^ CX_3_CR1^+^ effector-like, and Ly108^−^CX_3_CR1^−^ terminally exhausted cells, but to our surprise, Ly108^+^CX_3_CR1^+^ cells did not express Gfi1; a closer examination of progenitors found a prominent Gfi1^−^ fraction (tdTomato^lo^ cells). We contemplated that these cells may represent early progenitors differentiating to Gfi1^−^ Ly108^+^CX_3_CR1^+^ cells. To test this idea, considering that TCF1 is downregulated in differentiating progenitors, we analyzed TCF1 expression in tdTomato^lo^ cells, which showed a significant downregulation compared to Gfi1-tdTomato^hi^ cells (Fig. 1E). Altogether, we showed that Gfi1 expression is intricately linked to CD8 T cell exhaustion: first expressed on progenitors, decreased in a subset of progenitors and Ly108^+^CX_3_CR1^+^ cells, and then re-expressed in effector-like and terminally exhausted cells.

### Ly108^+^CX_3_CR1^+^ cells are a distinct subset with shared chromatin and transcriptional properties of progenitor and effector-like cells

Our analyses of Gfi1 expression suggest that Ly108^+^CX_3_CR1^+^ cells are a distinct population from other well-established subsets. We reason that Ly108^+^CX_3_CR1^+^ cells would possess distinct chromatin accessibility and transcriptomic profiles. To examine this, we sorted progenitor, Ly108^+^CX_3_CR1^+^, effector-like and terminally exhausted subsets of exhausted GP33^+^ cells from LCMV CL-13 infected mice on day 30 and performed ATAC-seq. The progenitor population is the source of the other subsets during chronic viral infection^(*6, 9*)^. We thus focused our initial analyses on identifying statistically significant chromatin changes with at least a 1.2 log2 fold change associated with the formation of Ly108^+^CX_3_CR1^+^ cells from progenitors. Ly108^+^CX_3_CR1^+^ cells had a chromatin profile with predominantly increased accessibility (Fig. 2A). Annotation of the identified differentially accessible chromatin regions (DACRs) showed that the majority of these changes were localized at distal intergenic and intronic regions (Fig. 2B) in accordance with the concept that enhancer regions define cellular identity^(*46*)^. Next, we visualized these DACRs to understand how they were organized in the effector-like and terminally exhausted subsets, which showed regions shared between Ly108^+^CX_3_CR1^+^ and effector-like cells with similar accessibility profiles (Fig. 2C-Box i and v and Fig. S2A); in other regions, the increase (Fig 2C, Box ii and S2A) or decrease (Fig. 2C-Box iv and Fig. S2A) in chromatin accessibility in Ly108^+^CX_3_CR1^+^ cells appeared intermediate to those in effector-like cells. Of note, several of the DACRs associated with Ly108^+^CX_3_CR1^+^ and effector-like cell formation were at loci with reported roles in T cell exhaustion and differentiation such as *Irf5*^(*47*)^, *Irf4*^(*48*)^, *Zeb2*^(*13*)^, *Batf3*^(*49*)^, *Batf*^(*9, 18, 50*)^, and *Hif1a*^(*51*)^ (Fig. S2A, with their corresponding Box numbers listed in parentheses). Finally, we also identified regions with increased accessibility specific to Ly108^+^CX_3_CR1^+^ cells (Fig. 2C-Box iii and Fig. S2A).

**Figure 2.**
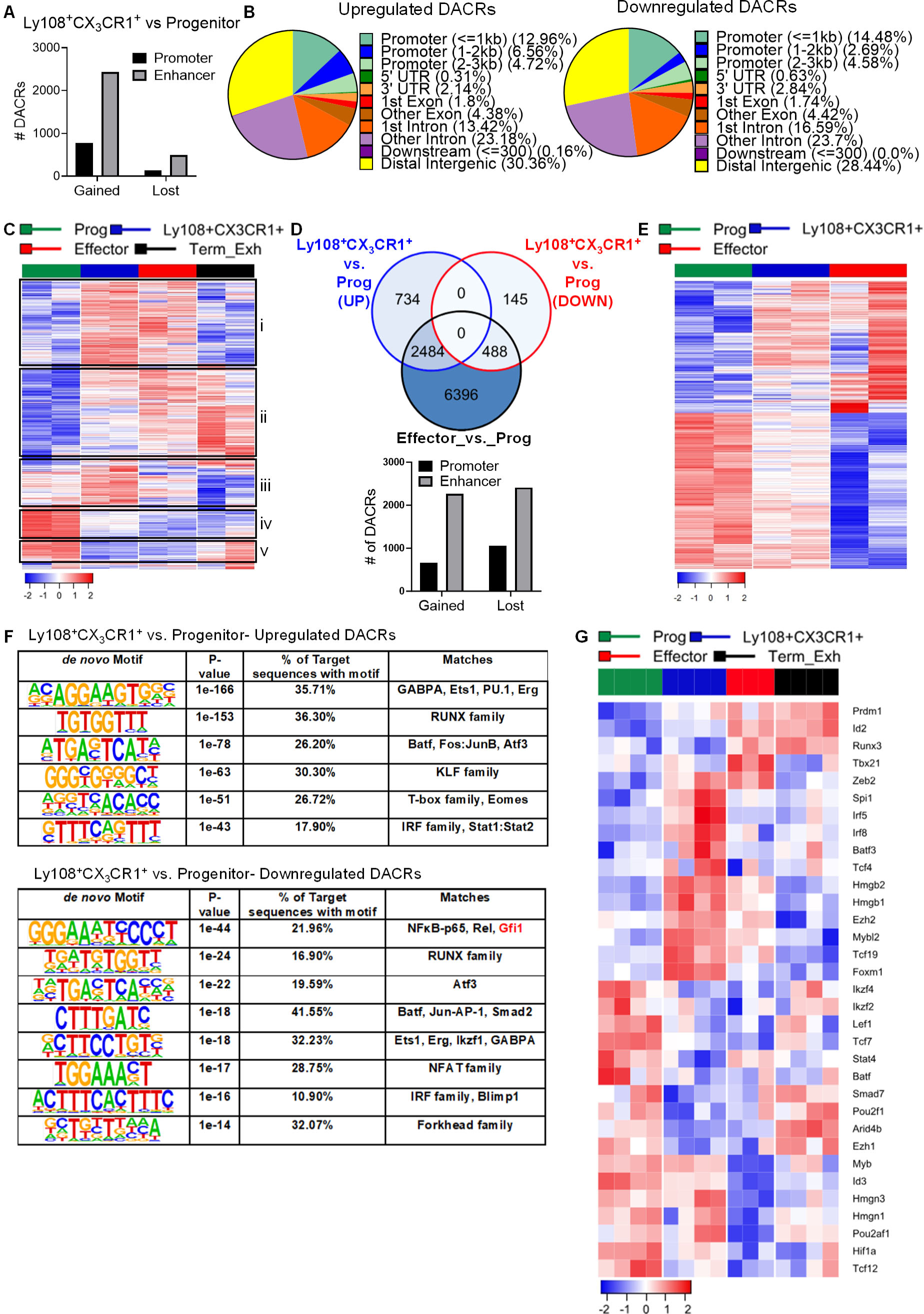
Ly108^+^CX_3_CR1^+^ cells are a distinct subset with shared chromatin and transcription properties of progenitor and effector-like cells. GP33^+^ CD8^+^ T cells were harvested and sorted from LCMV Cl-13 infected WT mice on day 30 based on Ly108 and CX_3_CR1 expression for ATAC-Seq (**A-F**) and RNA-Seq analyses (**G**). **A.** The number of differentially accessible chromatin regions (DACRs) in Ly108^+^CX_3_CR1^+^ cells versus progenitors (Ly108^+^CX_3_CR1^−^), with a cutoff of >1.2 log2 fold change. **B.** Upregulated (left) and downregulated (right) DACRs in Ly108^+^CX_3_CR1^+^ cells over progenitors. **C.** Heatmap visualization of DACRs in Ly108^+^CX_3_CR1^+^ cells versus progenitors in all WT subsets. Regions with distinct expression patterns among subsets are highlighted with roman numerals, Prog=Progenitor and Term_Exh=Terminally exhausted. **D.** Venn diagram depicting the intersection of DACRs (upregulated and downregulated) in Ly108^+^CX_3_CR1^+^ cells versus progenitors (Prog) with those in effectors (Eff) versus progenitors, with the summary of identified DACRs in effectors versus Ly108^+^CX_3_CR1^+^ cells shown by the bar graph. **E.** Heatmap visualization of all DACRs in Ly108^+^CX_3_CR1^+^ and effector cells over progenitors. **F.** HOMER motif analysis of DACRs. **H.** Heatmap visualization of select TFs among all the subsets. ATAC and RNA seq data were from 2 and 3-4 independent replicates, respectively. Each replicate was a pooled sample of 3-5 mice.

While 77% of identified DACRs between Ly108^+^CX_3_CR1^+^ and progenitors were maintained in effector-like cells (Fig. 2D, 2972 out of 3851 DACRs), 68% of DACRs between progenitors and effectors were unique to effector cells and not shared with Ly108^+^CX_3_CR1^+^ cells (6396 out of 9368 DACRs are unique to effector-like cells). Most of the effector-specific chromatin changes were also localized at distal intergenic and intronic regions (Fig. 2D). Using DACRs unique to effector cells, we found that Ly108^+^CX_3_CR1^+^ cells had an intermediate chromatin accessibility profile between progenitor and effector cells (Fig. 2E). The regions with increased accessibility in Ly108^+^CX_3_CR1^+^ and effector-like cells over progenitors were enriched for pathways involving GTPase activity (Fig. S2B) and downstream signaling leading to NFAT and AP1 activity that are important in T cell activation^(*52*)^. In contrast, regions with decreased accessibility were enriched for Wnt signaling known to inhibit effector differentiation^(*53*)^.

We next wanted to predict the transcription factors (TFs) interacting with the DACRs in the formation of Ly108^+^CX_3_CR1^+^ and effector-like cells by performing *de novo* motif analysis using HOMER^(*54*)^ (Fig. 2F and S2C). Motifs for TFs with previously reported roles in effector function such as Batf, AP-1/NFAT and IRF family proteins^(*11, 18, 47, 48, 50, 55–64*)^ were widely enriched in chromatin regions in both Ly108^+^CX_3_CR1^+^ and effector-like cells (Fig. 2F and S2C). In contrast, we observed that CTCF, a regulator of the 3D genomic architecture^(*65*)^ and linked to cytotoxic program genes (e.g.,*Tbx21*, *Ifng*, and *Klrg*^(*66*)^), was enriched only in regions involved in the formation of effector-like cells (Fig. S2C). In line with downregulated Gfi1 in Ly108^+^CX_3_CR1^+^ cells (Fig. 1D), the Gfi1 motif (5’-AAATC-3’) was enriched in downregulated DACRs between progenitor and Ly108^+^CX_3_CR1^+^ cells (Fig. 2F).

In support of these predicted TFs, bulk RNA-seq similarly identified *Batf*, *Irf5*, and *Zeb2* (Fig. 2G), and activation and effector molecules (Fig. S2D). Expression of *Tcf7* (and its surrogate marker *Slamf6*, Fig S2D) was consistent with its essential role in progenitors^(*4, 8, 11*)^, while *Zeb2*^(*13, 15*)^ expression was consistent with its involvement in the formation of effector cells. However, Irf8, previously reported to act independently of Tbet/Eomes and favor effector differentiation^(*63*)^, was selectively expressed in Ly108^+^CX_3_CR1^+^ cells. Recent studies described intermediate or precursor effector subsets based on Cxcr6^(*13*)^ and Klrg1^(*15*)^. However, Cxcr6 was not expressed in Ly108^+^CX_3_CR1^+^ cells (Fig. S2D-E), and Klrg1 was equally expressed in both Ly108^+^CX_3_CR1^+^ cells and Ly108^−^CX_3_CR1^+^ effector-like populations (Fig. S2F). Based on these data, the Ly108^+^CX_3_CR1^+^ cells present as a distinct subset, with unique as well as shared features to effector-like cells and progenitor cells; they are also different from previously described intermediate cells.

### CD8 T cell Gfi1 controls the formation of effectors and terminally exhausted cells

The dynamics of Gfi1 expression and the enrichment of the Gfi1 motif (5’-AAATC-3’) in DACRs of Ly108^+^CX_3_CR1^+^ cells prompted us to investigate whether Gfi1 regulates the development of T cell exhaustion. We employed genetic mice with specific deletion of Gfi1 in T cells (CD4-Cre x Gfi1^fl/fl(*35*)^) (hereafter Gfi1^cKO^) that we previously described^(*39, 40*)^. CD8 T cell responses were evaluated in Gfi1^fl/fl^CD4-Cre^−/−^ littermates (WT) and Gfi1^cKO^ mice between days 15-16 and 30-45, following LCMV-CL13 infection. Gfi1^cKO^ mice had lower numbers of total splenic CD8 as well as LCMV-GP33 and NP396 specific cells on day 15 (Fig. S3A). By day 30, the numbers of cells were comparable between Gfi1^cKO^ and WT mice (Fig. S3A). The frequencies of LCMV-specific cells did not differ at either time point (Fig. S3B). Next, we assessed whether Gfi1 deletion impacted the specific subsets of exhausted CD8 T cells. As shown in Fig. 3A (GP33^+^) and Fig. S3C (NP396^+^), Gfi1^cKO^ cells failed to produce effector and terminally exhausted cells and were arrested in the progenitor and Ly108^+^CX_3_CR1^+^ states, with consistently altered cell counts of GP33^+^ cells (Fig. S3D) and NP396^+^ cells (Fig. S3E). Failed formation of effector-like and terminally exhausted cells in Gfi1^cKO^ GP33^+^ CD8 T cells was also observed in the liver (Fig. S3F and S3G). Furthermore, Gfi1^cKO^ GP33^+^ (Fig. 3B) and NP396^+^ (Fig. S3H) cells did not express the effector molecule KLRG1. Altogether, Gfi1 deletion in T cells arrested the development of CD8 T cell exhaustion at the stage of Ly108^+^CX_3_CR1^+^, with no effector and terminally exhausted cells formed.

**Figure 3.**
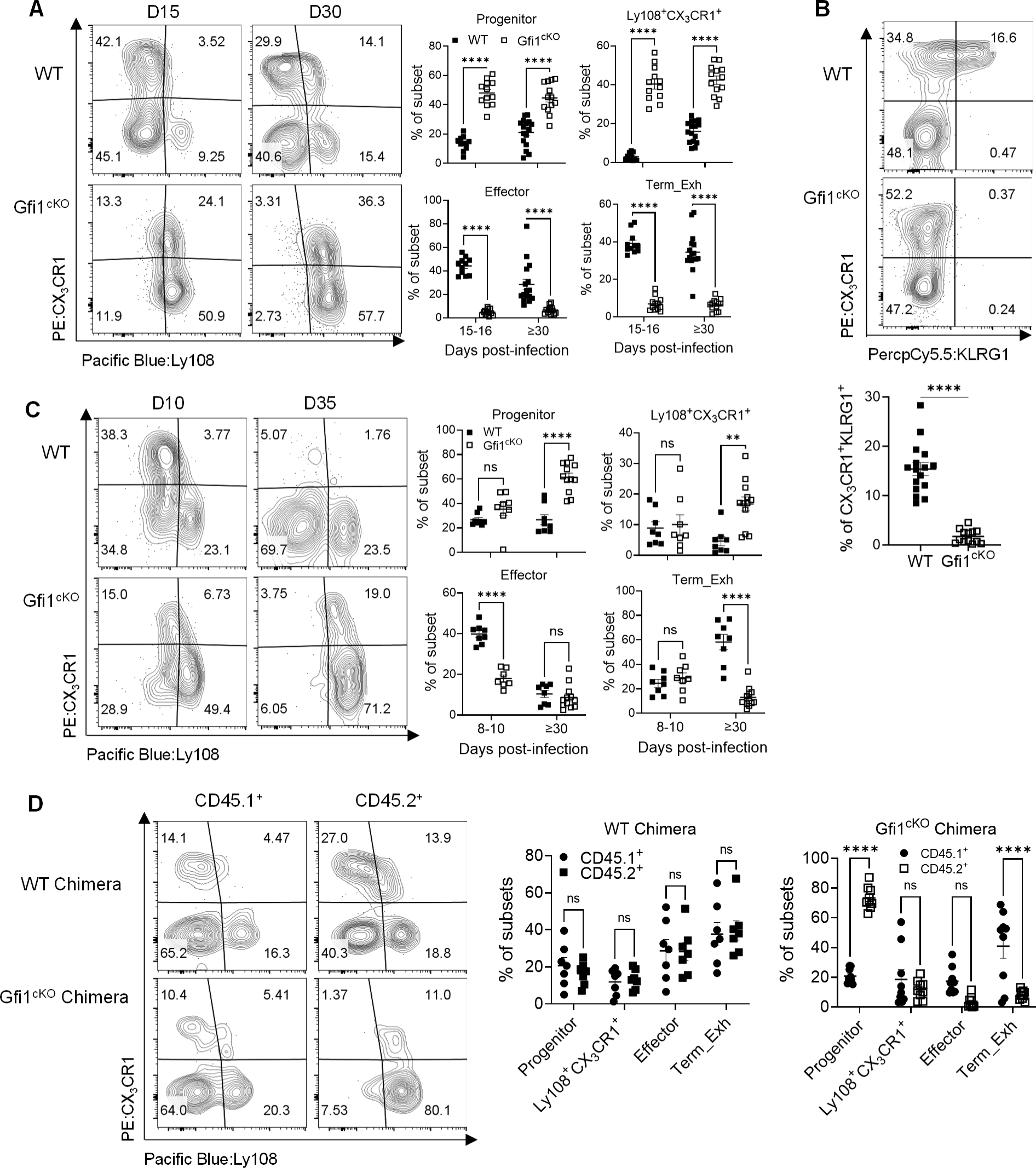
CD8 T cell-intrinsic Gfi1 controls the formation of effector and terminally exhausted cells. **A-B.** GP33^+^ CD8 cells from WT and Gfi1^cKO^ mice on day 15 and 30 post-LCMV Cl-13 infection were stained for CX_3_CR1 and Ly108 to determine the frequencies of the four subsets (**A**), and expression of CX_3_CR1 and KLRG1 on day 30 (**B)**. **C.** Mice depleted of CD4 T cells were infected with LCMV Cl-13, and at the designated times, GP33^+^ CD8 T cells were stained for CX_3_CR1 and Ly108 to determine the frequencies of the four subsets. **D.** WT/CD45.1 and Gfi1^cKO^ /CD45.1 chimeric mice were infected with LCMV Cl-13 for 30 days, and GP33^+^ CD8^+^ T cells within the CD45.1^+^ and CD45.2^+^ compartments were analyzed for CX_3_CR1 and Ly108. Data were from 3 independent experiments, shown as mean ± s.e.m., with each dot denoting an individual mouse. ns, P≥0.05; **, P<0.01; ***, P<0.001; ****, P<0.0001. Data in **A-C** were analyzed by two-way ANOVA with Sidak’s post-hoc test and in **D** by unpaired two-tailed Student’s t test.

Since CD4 T cell deficiency ablates the formation of effector-like exhausted CD8 T cells^(*16, 67*)^ and because Gfi1 is deleted in both CD8 and CD4 T cells in Gfi1^cKO^ mice, we wanted to rule out the potential confounding effects from Gfi1^cKO^ CD4 T cells. Depletion of CD4 T cells promoted exhaustion in both GP33^+^ (Fig. 3C versus 3A) and NP396^+^ (Fig. S3I versus S3C) WT CD8 T cells, and as expected, resulted in the absence of effector-like cells at day 30. Nevertheless, this did not change the phenotype of GP33^+^ and NP396^+^ Gfi1^cKO^ cells, as they remained arrested in progenitor and Ly108^+^CX_3_CR1^+^ states (Fig. S3J), suggesting this was likely a CD8 T cell-intrinsic effect. To firmly establish this, we used bone marrow chimeras reconstituted with either a mixture of congenically marked wild-type cells or wild-type cells with Gfi1^cKO^ bone marrow cells. Following reconstitution, these mice were infected with LCMV CL-13, and on day 30, CD8 T cells were analyzed. While in CD45.1^+^/CD45.2^+^ WT chimeras both CD45.1^+^ and CD45.2^+^ CD8 T cells successfully generated progenitor, Ly108^+^CX_3_CR1^+^, effector-like, and terminally exhausted GP33^+^ (Fig. 3D) and NP396^+^ CD8 T cells (Fig. S3K), in contrast, CD45.2^+^ Gfi1^cKO^ CD8 T cells in the CD45.1^+^/CD45.2^+^ Gfi1^cKO^ chimeras could not form effector-like and terminally exhausted GP33^+^ and NP396^+^ CD8 T cells (Fig. 3D and S3K). These data highlight a cell-autonomous role of Gfi1 in sustaining the normal differentiation of exhausted CD8 T cells, and Gfi1 deletion in T cells severely impair the formation of effector and terminally exhausted populations.

### Gfi1 maintains the chromatin organization and transcriptomic programs for the formation of exhausted effector CD8 T cells

Exhausted CD8 T cells subsets have distinct transcriptional and epigenetic profiles (Fig. 2)^(*6, 15, 18–21*)^. A recent study showed that epigenetic mechanisms govern the formation of T cell subsets during chronic viral infection^(*68*)^. Considering the differential expression of Gfi1 in the progenitor, Ly108^+^CX_3_CR1^+^, effector-like, and terminally exhausted subsets (Fig. 1D-E) and its reported role in chromatin modifying complexes in other cellular contexts^(*32–34*)^, we wanted to evaluate whether Gfi1 impacts the chromatin accessibility in exhausted CD8 T cells during chronic viral infection, which has not been explored before. To address this, we sorted LCMV GP33^+^ progenitor and Ly108^+^CX_3_CR1^+^ cells from LCMV CL-13 infected WT and Gfi1^cKO^ mice on day 30 post-infection. We also sorted WT effector-like cells as control for our ATAC-seq and RNA-seq analyses. Gfi1^cKO^ progenitor and Ly108^+^CX_3_CR1^+^ cells exhibited considerable changes in chromatin accessibility compared to their WT counterparts (Fig. 4A). Most of these changes were localized at distal intergenic and intronic regions with few at promoter regions (Fig. 4B and S4A-C). In keeping with Gfi1 being a transcriptional repressor, Gfi1^cKO^ progenitors had a significant increase in chromatin accessibility compared to WT progenitors, many of which were restricted for WT Ly108^+^CX_3_CR1^+^ and effector-like cells (Fig. 4C, Box ii). Given this, we asked if the normal chromatin reorganization during the differentiation of progenitors to Ly108^+^CX_3_CR1^+^ cells was disrupted by Gfi1 deletion. Strikingly, very few DACRs (only 30 upregulated and 164 downregulated) were identified in Gfi1^cKO^ Ly108^+^CX_3_CR1^+^ cells versus progenitors, in contrast to 3218 upregulated and 633 downregulated DACRs in WT cells (Fig. 4D and S4D). Understandably, some of the chromatin reorganization in WT Ly108^+^CX_3_CR1^+^ cells that gained (Fig. 4C, Box i) or lost accessibility (Fig. 4C, Box iii) was retained in Gfi1^cKO^ Ly108^+^CX_3_CR1^+^ cells.

**Figure 4.**
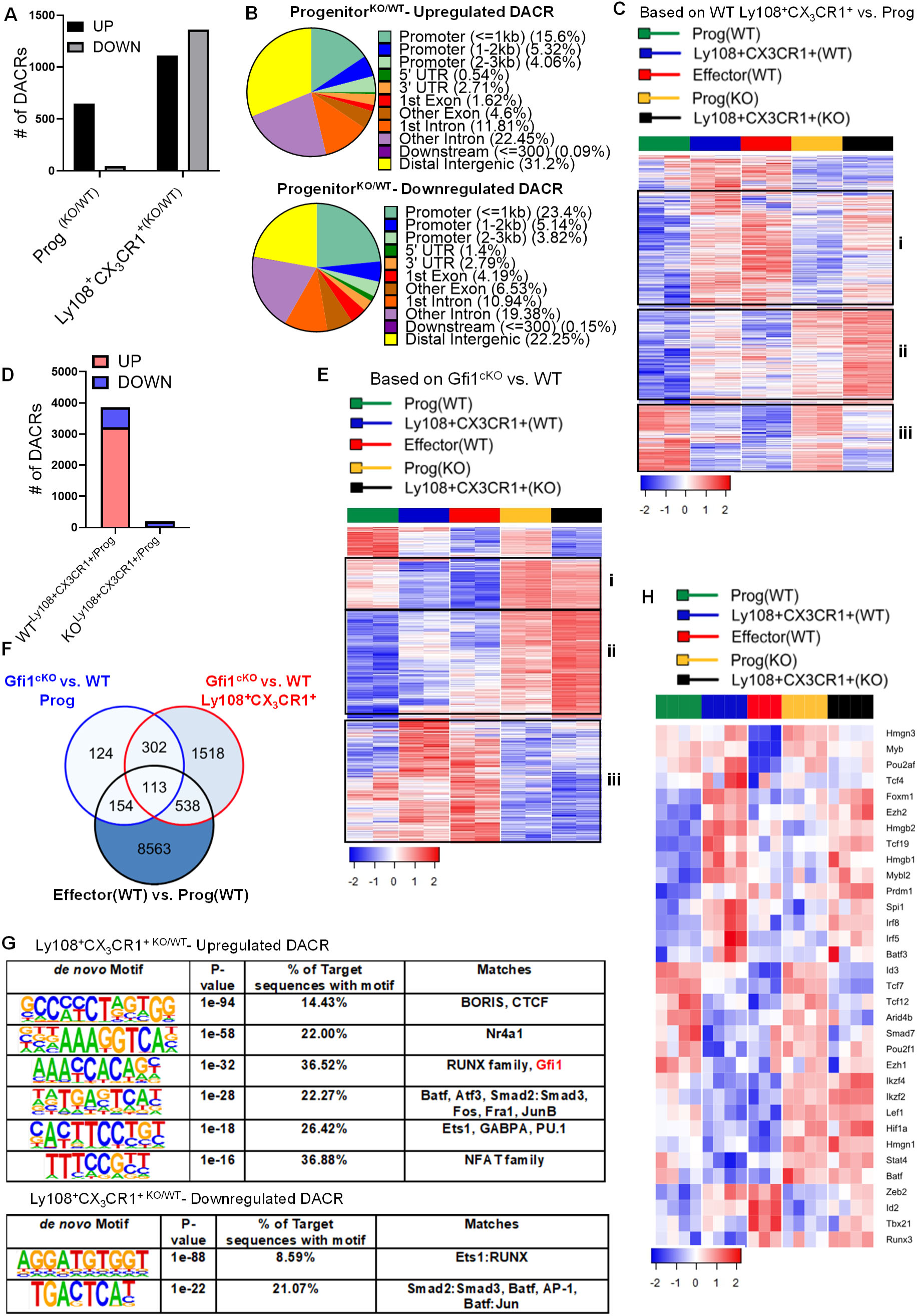
Gfi1 maintains the chromatin accessibility needed for the formation of effector cells. GP33^+^ exhausted CD8^+^ T cell subsets were harvested and sorted from LCMV Cl-13 infected WT and Gfi1^cKO^ mice on day 30 for ATAC-seq or RNA-seq. **A.** Differentially accessible chromatin regions (DACRs) between Gfi1^cKO^ versus WT progenitors as well as between Gfi1^cKO^ versus WT Ly108^+^CX_3_CR1^+^ cells, with a cutoff of 1.2 log2 fold change. **B.** Distribution of upregulated (upper) and downregulated (lower) DACRs between Gfi1^cKO^ and WT progenitors. **C.** Heatmap visualization of DACRs in WT Ly108^+^CX_3_CR1^+^ cells over progenitors among all the WT and Gfi1^cKO^ subsets. DACRs with distinct expression patterns were highlighted with Roman numerals, Prog=Progenitor, Eff=Effector, and Term_Exh=Terminally Exhausted. **C.** DACRs between WT Ly108^+^CX_3_CR1^+^ cells versus progenitors and between Gfi1^cKO^ Ly108^+^CX_3_CR1^+^ cells versus progenitors. **E.** Heatmap visualization of total DACRs in Gfi1^cKO^ Ly108^+^CX_3_CR1^+^ cells and progenitors over their WT counterparts. DACRs with distinct expression patterns were highlighted with Roman numerals. **F.** Intersection of the DACRs between WT effectors (Eff) versus progenitors (Prog), Gfi1^cKO^ versus WT Ly108^+^CX_3_CR1^+^, and Gfi1^cKO^ versus WT progenitors. **G.** HOMER motif analysis of identified DACRs. **H.** Heatmap visualization of transcripts for select TFs in WT and Gfi1^cKO^ GP33^+^ subsets. ATAC-seq and RNA-seq data were from 2 and 3-4 independent replicates, respectively. Each replicate was a pooled sample from 3-5 mice.

To determine how Gfi1 loss caused the significant reduction of effector-like cells, we focused on the total DACRs in both Gfi1^cKO^ progenitor and Ly108^+^CX_3_CR1^+^ cells versus corresponding WT subsets. Heatmap visualization revealed significant dysregulation in DACRs associated with effector-like cells (Fig. 4E). Specifically, some regions failed to lose (Box i in Fig. 4E) or gain (Box iii in Fig. 4E) accessibility in Gfi1^cKO^ cells. Examples of the regions impacted by Gfi1 deficiency were shown in Fig. S4E, including *Lef1*^(*69*)^, *Irf5*^(*47*)^*, Batf3*^(*49*)^, and *Mef2c*^(*70*)^, which have been implicated in orchestrating genomic architecture and T cell effector functions. There were also DACRs with greatly increased accessibility in Gfi1^cKO^ cells, regardless of subset identity, as compared to their WT counterparts (Box ii in Fig. 4E), such as regions in *Snai2* and *Ezh1* (Fig. S4E). Of note, Ezh1 is an integral component of the polycomb repressive complex, and its repression facilitates the formation of effector and memory T cell subsets^(*71*)^. GO enrichment analysis of upregulated DACRs in Gfi1^cKO^ Ly108^+^CX_3_CR1^+^ cells revealed pathways involving GTPase activity, DNA repression activity, and HMG box domain activity (Fig.S4F), with reported role in T cell biology^(*52*)^. To understand potential TFs mediating the chromatin dysregulation in Gfi1^cKO^ cells, we performed *de novo* motif analysis using the DACRs between WT and Gfi1^cKO^ cells and found enrichment of motifs for differentiation regulators such as Batf^(*18, 50*)^, CTCF^(*65*)^, and NFAT (Fig. 4G and S4G). Moreover, in support of the auto-repression of *Gfi1* locus by Gfi1 itself^(*31*)^, Gfi1 motifs were enriched in accessible regions in Gfi1^cKO^ cells (Fig. S4G and 4G). Noteworthily, the chromatin changes in Gfi1^cKO^ CD8 T cells were only seen in antigen specific exhausted cells, as the chromatin profiles in naïve Gfi1^cKO^ cells appeared normal (Fig. S4H).

To understand how the dysregulated chromatin architecture in Gfi1^cKO^ cells affected gene expression, we performed RNA-seq. Consistent with the maintenance of some chromatin structures and the formation of progenitor and Ly108^+^CX_3_CR1^+^ cells in Gfi1^cKO^ mice, transcripts for some subset specific TFs (*Tcf7*, *Spi1*, and *Irf8*) were largely unaltered in Gfi1^cKO^ cells (Fig. 4H). In contrast, transcripts for *Batf*, *Lef1*, *Ezh1*, *Ikzf2*, *Stat4*, and *Hmgn1* that were normally downregulated in WT effector-like and Ly108^+^CX_3_CR1^+^ cells, remained highly expressed in Gfi1^cKO^ Ly108^+^CX_3_CR1^+^ cells. Similarly, transcripts for activation and adhesion molecules, including *Klrg1, Itgax, Havcr2, Tigit, Lag3*, and *Glp1r*, were also dysregulated in Gfi1^cKO^ cells (Fig. S4I). Overall, these data indicate that Gfi1 repressed several key differentiation programs that were necessary for the chromatin rewiring needed for exhausted effector-like cell formation.

### Gfi1 is required for the generation of terminally differentiated intratumoral CD8 T cells

Since our findings demonstrated that Gfi1 regulated the generation of effector-like and terminally exhausted cells during chronic LCMV CL-13 infection, we asked if Gfi1 was also required for the formation of terminally differentiated effector tumor infiltrating CD8 T cells (CD8 TILs) known to be exhausted. To explore this, Gfi1^tdTomato^ reporter mice bearing MB49 urothelial adenocarcinoma cells were euthanized at various stages of tumor progression to analyze CD8 TILs. CD8 TILs can be largely grouped into TCF1^+^ progenitors and terminally differentiated Tim3^+^ effector cells^(*7, 72*)^, with the latter producing some effector molecules including perforin and granzyme to exert some control of tumor growth. As shown in Fig. 5A, while TCF1^+^ progenitor cells were the dominant subset of CD8 TILs during the early stage, Tim3^+^ terminally differentiated effector cells became dominant during the later stages of tumor progression. Further characterization of total CD8 TILs based on tdTomato (Gfi1) expression revealed dynamic Gfi1 expression during tumor progression (Fig. S5A). We also found that TCF1^hi^ progenitor TILs expressed higher levels of Gfi1 compared to Tim3^hi^ cells (Fig. 5B). Tumor progression was associated with a decrease in tdTomato^lo^ progenitor TILs (Fig. S5B), which appeared to be the differentiating progenitor TILs, based on the downregulation of TCF1 (Fig. 5C), as in our LCMV studies (Fig. 1E).

**Figure 5.**
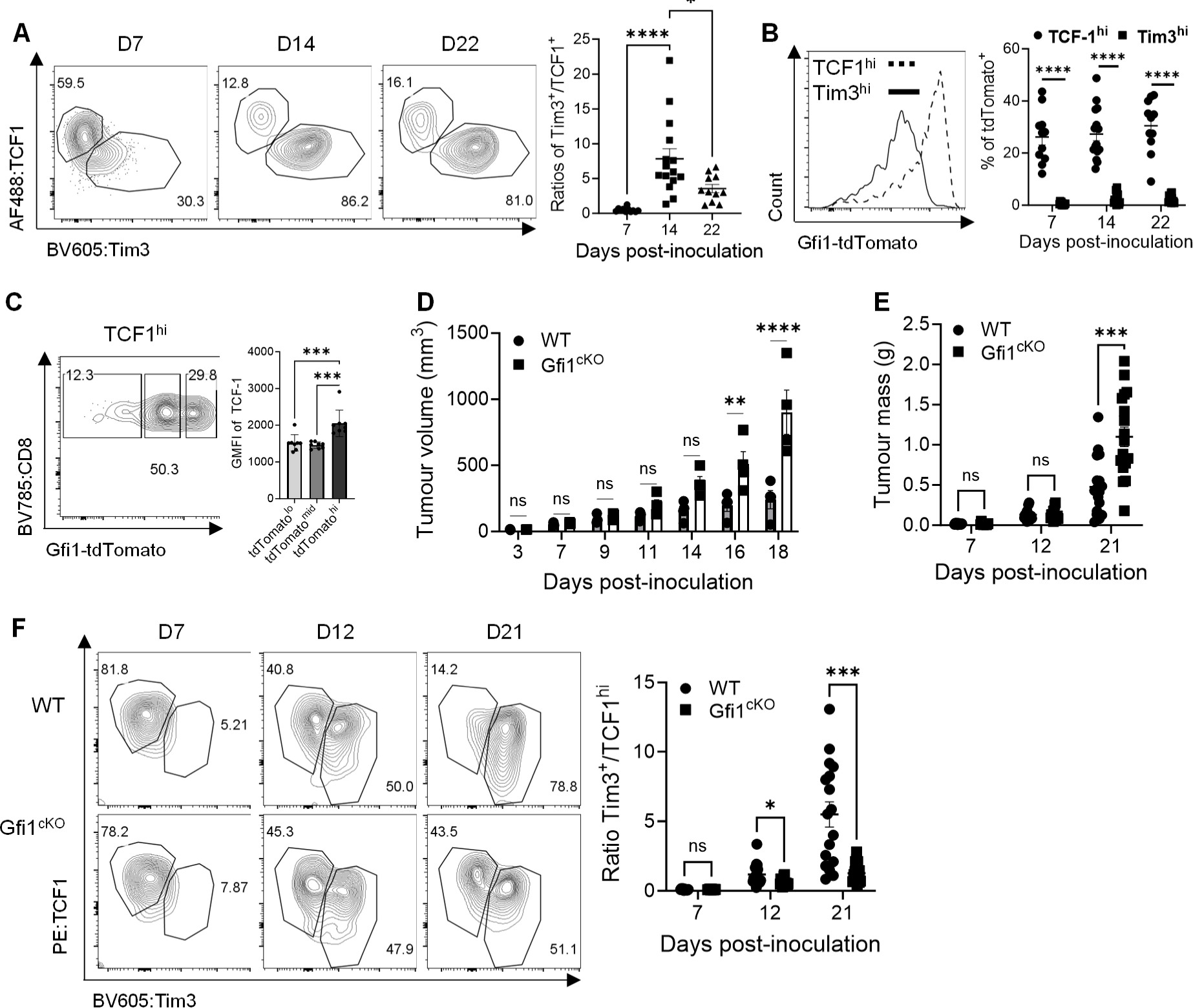
Gfi1 is required for the generation of terminally differentiated intratumoral CD8^+^ T cells. **A-C.** Gfi1^tdTomato^ reporter mice were inoculated with MB49 urothelial adenocarcinoma cells. At different times post-tumor inoculation, tumor infiltrating CD8^+^ T cells (CD8^+^ TILs) were stained for TCF1 and Tim3 to determine progenitors (TCF1^hi/+^) and terminally differentiated effector cells (Tim3^hi/+^), with ratios of Tim3 ^hi^/TCF1^hi^ cells shown by the scatter dot plot (**A**), tdTomato/Gfi1 expression in TCF1^hi^ and Tim3^hi^ cells (**B**), and GMFIs of TCF1 in tdTomato^lo^, tdTomato^mid^, and tdTomato^hi^ subsets among TCF1^+^ cells (**C**). **D-F.** WT and Gfi1^cKO^ mice bearing MB49 tumors were closely monitored for tumor growth (**D**) and at the designated times, sets of mice were euthanized to measure tumor weights (**E**); isolated CD8^+^ TILs from these mice were stained for TCF1 and Tim3, with ratios of Tim3^+^/TCF1^+^ CD8^+^ TILs shown in the scatter dot plot (**F**). Data were pooled results from 2-4 independent experiments, shown as mean ± s.e.m., with each dot denoting an individual mouse. ns, P≥0.05; *, P<0.05; **, P<0.01; ***, P<0.001; ****, P<0.0001. Data in **A,C** were analyzed by one-way ANOVA with Tukey’s or Dunnett’s post-hoc test, and data in **B**,**D-F** were analyzed by two-way ANOVA with Sidak’s post hoc test.

Next, we assessed how Gfi1 deletion in T cells impacted the transition of TCF1^+^ progenitor TILs to terminally differentiated Tim3^+^ effector TILs. To this end, WT and Gfi1^cKO^ mice were inoculated with MB49 tumor cells, and tumor growth was regularly measured. While Gfi1 was seemingly dispensable for tumor control during the early phase (prior to day 14), Gfi1^cKO^ mice were unable to control tumor as effectively as WT mice beyond 16 days (Fig. 5D), consistent with our previous study showing that Gfi1^cKO^ mice could not control the B16 melanoma growth at the late stage^(*40*)^. Together, these results support an important role of T cell Gfi1 in controlling tumor progression. Tumor weights in Gfi1^cKO^ mice were also significantly greater than those in WT mice on day 21 but comparable on day 7 and day 12 (Fig. 5E). While our analyses of isolated TILs at these different stages of tumor growth did not reveal overt changes of CD4 TILs, evidenced by comparable frequencies of total CD4 TILs (Fig. S5C) and regulatory T cells (T_reg_) (Fig. S5D) in Gfi1^cKO^ versus WT mice, CD8 T cell infiltration in Gfi1^cKO^ mice increased on day 7 but then reduced on day 12 and 21 (Fig. S5C). Furthermore, Gfi1^cKO^ CD8 T cells were impaired to generate terminally differentiated effector cells at later time points (Fig. 5F and S5E), similar to what we found in the LCMV studies. These data indicate that Gfi1 is also important for CD8 TILs, by facilitating the transition of progenitors to terminally differentiated cells during tumor progression.

### Gfi1 expression in T cells is required for anti-CTLA-4 response

Our above results implicate a role of CD8 T cell Gfi1 activity in the transition of TCF1^+^ progenitors to terminally differentiated Tim3^+^ effector TILs during tumor progression. Considering that ICBs such as anti-CTLA-4/PD-1 rely on this transition to mediate their therapeutic effects^(*7, 8, 12*)^, we contemplated that Gfi1 expression in T cells would be needed for ICB efficacy. Using anti-CTLA-4 as a prototypical ICB, we first asked if it would regulate Gfi1 expression in CD8 TILs. We analyzed TILs from MB49 tumor-bearing Gfi1^tdTomato^ mice, treated with or without anti-CTLA-4. As expected^(*73*)^, anti-CTLA-4 significantly increased infiltration of both CD4 and CD8 T cells (Fig. S6A). Importantly, anti-CTLA-4 significantly reduced the frequencies of Gfi1-tdTomato^hi^ CD8 progenitor TILs (Fig. S6B), indicating an active regulation of Gfi1 expression in CD8 TILs by ICBs. To determine if Gfi1 in T cells mediates therapeutic effects of anti-CTLA-4, WT and Gfi1^cKO^ mice bearing established MB49 tumors were treated with anti-CTLA-4^(*73*)^. Consistent with our previous study^(*73*)^, anti-CTLA-4 potently suppressed tumor growth in WT mice (Fig. 6A) but was unable to do so in Gfi1^cKO^ mice (Fig. 6B), indicating an essential role of T cell Gfi1 in anti-CTLA-4. This was accompanied by significantly reduced IFN-γ production in Gfi1^cKO^ CD8 TILs (Fig. 6C), although functional rejuvenation of CD4 TILs and T_reg_ depletion by anti-CTLA-4 seemed unaltered by Gfi1 loss (Fig. S6C-D). To explore this in a different tumor model, we inoculated WT and Gfi1^cKO^ mice with MC38 colorectal tumor cells. Like MB49 bladder tumors, anti-CTLA-4 suppressed MC38 colorectal tumors in WT but not Gfi1^cKO^ mice (Fig. 6D). Unlike MB49 bladder tumor, anti-CTLA-4 did not induce CD4 T cell infiltration in MC38 colorectal cancer (Fig. 6E) even in WT mice. Nevertheless, augmentation of IFN-γ production (Fig. S6E) and depletion of T_reg_ (Fig. S6F) by anti-CTLA-4 persisted in both WT and Gfi1^cKO^ tumor-bearing mice, further supporting a largely dispensable role of Gfi1 in CD4 TILs. With respect to CD8 TILs, while anti-CTLA-4 promoted CD8 T cell infiltration in both WT and Gfi1^cKO^ MC38 tumor-bearing mice (Fig. 6E), the effect was much weaker in the latter. Moreover, anti-CTLA-4 only increased IFN-γ production in WT but not Gfi1^cKO^ CD8 TILs (Fig. 6F). In summary, Gfi1 deletion in T cells rendered MB49 and MC38 tumor-bearing mice non-responsive to anti-CTLA-4, coupled with primary defects in CD8 but not CD4 TILs, highlighting Gfi1 loss in CD8 T cells as a major mechanism of therapeutic resistance to ICBs. These results corroborated the CD8 T cell-autonomous role of Gfi1 in our LCMV studies (Fig. 3), that is, the impaired formation of exhausted effector-like CD8 T cells in the absence of Gfi1.

**Figure 6.**
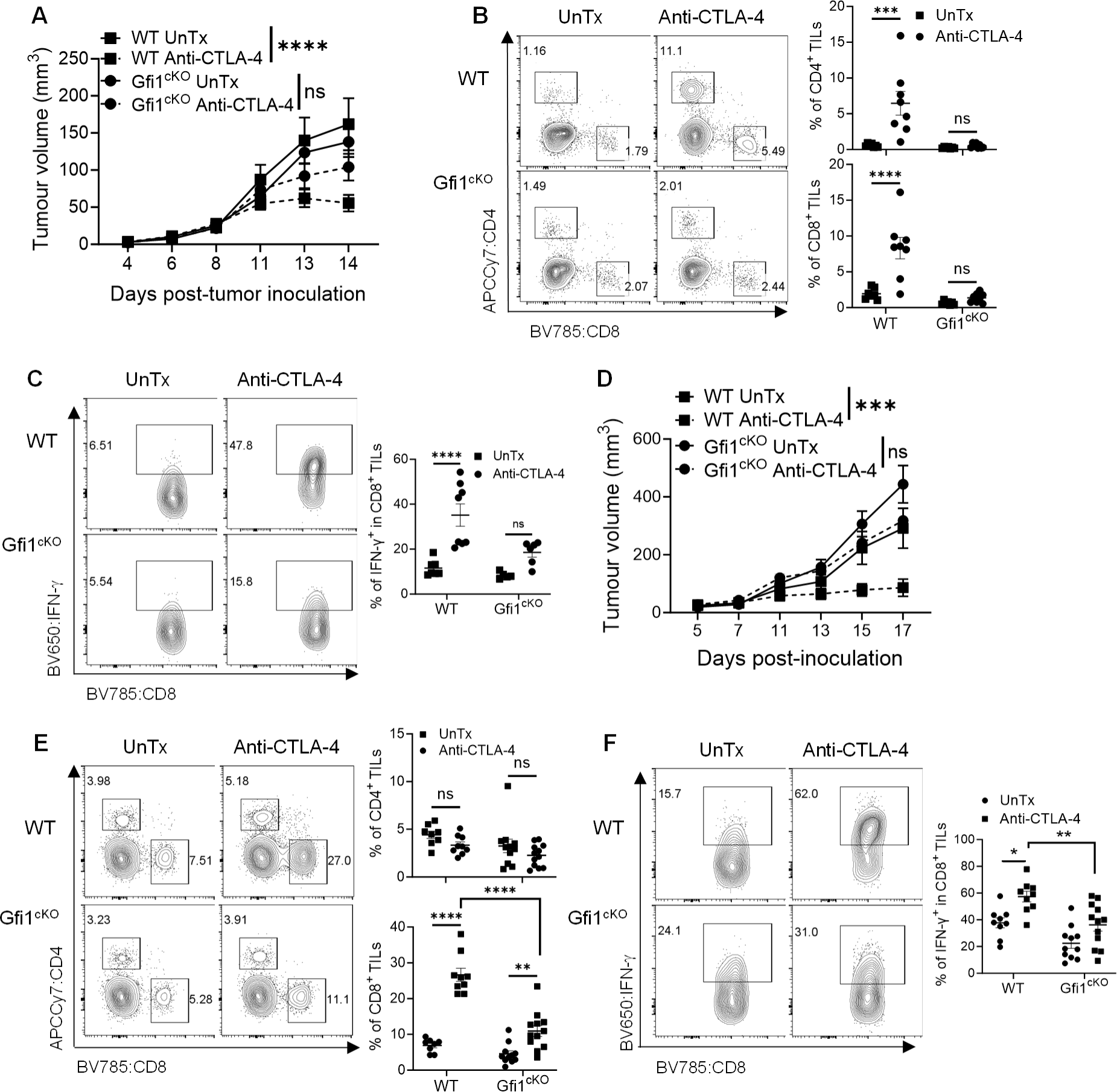
T cell-intrinsic Gfi1 controls anti-CTLA-4 therapeutic response. WT and Gfi1^cKO^ mice bearing palpable MB49 bladder tumor (**A-C**) or MC38 colorectal tumor (**D-F**) were treated with anti-CTLA-4, followed by periodic measurements of tumor growth (**A, D**). Mice were then euthanized to determine the infiltration of CD4^+^ and CD8^+^ T cells (**B, E**) and IFN-γ production by CD8^+^ TILs (**C, F**). Data were pooled results of 2 independent experiments, shown as mean ± s.e.m., with each dot denoting an individual mouse. ns, P≥0.05; *, P<0.05; **, P<0.01; ***, P<0.001; ****, P<0.0001. Data were analyzed by two-way ANOVA with Sidak’s post hoc test.

## Discussion

We have identified a new role of Gfi1 in CD8 T cell exhaustion. Gfi1 expression is dynamically regulated during chronic LCMV CL-13 infection in that it is not expressed during the early stage, emerges first in progenitor cells on days 12-15 when T cell exhaustion begins to develop^(*9*)^ and then on effector-like and terminally exhausted cells by day 30 when T cell exhaustion is established, but it is downregulated in a fraction of progenitor cells as well as Ly108^+^CX_3_CR1^+^ cells. This dynamic induction and regulation of Gfi1 expression during T cell exhaustion supports the model proposed by Utzschneider et al.^(*9*)^ that early LCMV-specific effectors are replaced by dysfunctional effector-like and terminally exhausted cells generated from exhausted progenitor populations at later stages of chronic viral infection. Of note, unlike the phenotypes from the perturbation of other well-defined TFs in CD8 T cell exhaustion (e.g., TOX, TCF1, Batf, Zeb2^(*9, 13, 22, 25*)^, Gfi1 deletion in T cells leads to a unique phenotype, that is, lack of effector-like and terminally exhausted CD8 T cells with accumulation of progenitors and Ly108^+^CX_3_CR1^+^ cells. The Ly108^+^CX_3_CR1^+^ cells appear to be a distinct subset with unique chromatin accessibility profile and transcriptome as well as shared features with progenitors and effector-like cells. Gfi1 deletion disrupts the chromatin remodeling and transcriptional programs in progenitors and Ly108^+^CX_3_CR1^+^ cells needed for the formation of effector-like and terminally exhausted cells. Similarly, TILs lacking Gfi1 are also impaired to transit from TCF1^hi^ progenitors to terminally differentiated Tim3^hi^ effector TILs. Consequently, Gfi1^cKO^ tumor-bearing mice are less able to control tumor progression and not responsive to ICB anti-CTLA-4. Our results highlight Gfi1 as a critical TF in CD8 T cell exhaustion, which may offer a new avenue to fine-tune this process.

Current models of CD8 T cell exhaustion propose that self-renewing exhausted progenitors differentiate into either effector-like or terminally exhausted fates^(*13–15*)^. Our analyses of the chromatin accessibility (Fig 2E) suggest that Ly108^+^CX_3_CR1^+^ cells may add another component to these models, with a potential to further mature into Ly108^−^CX_3_CR1^+^ effector-like cells. We also identify a diversification of progenitors to Gfi1^+^ and Gfi1-fractions. Further elaboration on whether Gfi1^+^ progenitors give rise to effector-like and terminally exhausted subsets while Gfi1^−^ progenitors form Ly108^+^CX_3_CR1^+^ cells will further enrich these models, warranting future investigations. Specific deletion of Gfi1 in T cells disrupts the normal chromatin landscape and transcriptome in progenitor and Ly108^+^CX_3_CR1^+^ cells, many of which are essential for the appropriate genomic architecture and the development of exhausted CD8 T cell subsets. Most subset specific DACRs occur at distal intergenic and intronic regions, which are known to define cellular identity^(*46*)^. CTCF and its co-factor BORIS in DACRs are selectively enriched in WT and Gfi1^cKO^ Ly108^+^CX_3_CR1^+^ cells but not progenitors. It will be interesting to investigate their functional interactions with Gfi1. Along the same line, given the well-established functional antagonism between PU.1 and Gfi1^(*54, 74–76*)^ and selective expression of PU.1 in Ly108^+^CX_3_CR1^+^ cells, future endeavors to examine how PU.1 regulates exhausted CD8 T cell subsets will be informative, as there has not been studied.

Similar to the inability of Gfi1^cKO^ LCMV-specific progenitors to form exhausted effector-like CD8 T cells, Gfi1^cKO^ progenitor TILs are also impaired in their ability to generate terminally differentiated effector TILs at later time points. Although our tumor data largely corroborate our LCMV CL-13 results, there are some gradient discrepancies between them, which could be simply due to the fundamental differences between chronic infections and cancer, as viral antigens are exogenous and highly immunogenic, whereas tumor antigens are mostly endogenous molecules with some degree of mutation and weakly immunogenic. Alternatively, it is possible Tim3 underestimates the heterogeneity of terminally differentiated effector cells of TILs. Nevertheless, our results show that Gfi1 activity in T cells regulates the formation of exhausted effector cells in both settings. Despite the unprecedented clinical successes of ICBs (anti-CTLA-4, anti-PD-1/L1, anti-LAG3, etc.) in various types of advanced cancer, therapeutic resistance to ICBs has emerged as a pressing issue. We identify loss of Gfi1 in CD8 T cells as a major T cell-intrinsic mechanism of therapeutic resistance to ICBs. Because the development of T cell exhaustion is a major barrier to effective immunotherapeutic approaches (e.g. CAR-T and ICBs)^(*4, 8, 77, 78*)^, identifying TFs and signals that regulate Gfi1 protein expression in progenitors will allow for the manipulation of T cell exhaustion to benefit patients. In conclusion, we show that the transcriptional repressor Gfi1 is an important regulator of the development of exhausted CD8 T cells during both chronic infections and tumors.

## Methods

### Mice

Gfi1tdTomato mice were generated by Dr. George Lacaud^(*43*)^ and transferred to us from Dr. Lee Grimes, with permission from Dr. Lacaud. Gfi1^fl/fl^CD4^Cre^ mice, as we previously described^(*39, 40*)^, were generated by Dr. Hano Hock^(*35*)^ and provided to us by Dr. Hongbo Chi, in agreement with Dr. Hock. Male and female Gfi1^fl/fl^CD4^Cre^ were bred as heterozygotes at the CD4-Cre locus (Gfi1^cKO^) and littermates (CD4-Cre^−/−^) were used in all experiments as wild-type controls (WT). B6.SJL-PtprcaPepcb/BoyJ mice (CD45.1) were obtained from the Jackson Laboratory. CD45.1 mice were crossed with Gfi1^fl/fl^CD4-Cre^−/−^ (WT) mice to generate CD45.1^+^CD45.2^+^ mice, which were used as recipients in the bone marrow chimera studies. All mice were bred and maintained in specific pathogen-free conditions in the animal facility of The University of Alabama at Birmingham (UAB) under 12 hours/12 hours light/dark cycle, ambient room temperature (22 °C) with 40-70% humidity. Seven-twelve weeks old mice were used in the experiments, and protocols were approved by Institutional Animal Care and Use Committee at UAB.

### LCMV Infection challenges, bone marrow chimeras

For chronic infections, 4×10^6^ plaque-forming-units (PFU) of LCMV CL-13 were intravenously injected into Gfi1^cKO^ and littermate control mice via tail vein. To generate the mixed bone marrow chimeras, recipient CD45.1^+^CD45.2^+^ mice were lethally irradiated with 2 doses of 500 Rad with an at least 4-hour interval between exposures and reconstituted with 1×10^7^ bone marrow cells depleted of mature T cells from CD45.1^+^ and CD45.2^+^ Gfi1^fl/fl^CD4Cre^−/−^ (WT) or Gfi1^fl/fl^CD4Cre^+/−^ (Gfi1^cKO^) mixed at 1:1 ratio. After at least 6 weeks of reconstitution, mice were used in the LCMV CL-13 infection experiments. In the CD4 T cell depletion experiments, *in vivo* neutralizing antibodies against CD4 (Bio X Cell, clone GK1.5, #BE0003-1) (300 μg/mouse) were injected intraperitoneally 1 day before and 3 days after LCMV infection.

### In vivo tumor inoculation and treatment

The chemically induced murine urothelial adenocarcinoma MB49 cells and colon adenocarcinoma MC38 cells, kindly provided by Dr. Ashish Kamat and Dr. Jim Allison at MD Anderson Cancer Center, was originally derived from a male and a female C57BL/6 mouse, respectively. These cells were cultured in Dulbecco’s modified Eagle’s medium with 10% fetal bovine serum (FBS) at 37 °C, 5% CO_2_, with regular testing for mycoplasma using the MycoAlert detection kit (Lonza, LT07-118) to ensure they were free of mycoplasma contamination. For tumor inoculation, exponentially growing MB49 cells (1×10^5^) and MC38 cells (5×10^5^) with a passage number less than 10 were subcutaneously injected into the shaved right flanks of mice. On day 5 post-tumor inoculation, mice received anti-CTLA-4 or isotype antibodies intraperitoneally (i.p.), with the first dose at 200 μg/mouse and subsequent injections of 100 μg/mouse on day 8 and 11.Tumor measurements were taken 2–3 times/week, starting from day 3 post-tumor inoculation. Mice were euthanized when tumor reached 2.0 cm in diameter, ulceration occurred, or mice became moribund.

### TIL isolation and tissue preparation

Detailed procedures were described previously^(*79, 80*)^. In brief, tumors were collected in ice-cold RPMI 1640 containing 2% FBS and minced into fine pieces, followed by digestion with 400 U/mL collagenase D (Worthington Biochemical Corporation, #LS004186) and 20 µg/mL DNase I (Sigma, #10104159001) at 37 °C for 40 min with periodic shaking. EDTA (Sigma, #1233508) was then added at the final concentration of 10 mM to stop digestion. Cell suspensions were filtered through 70 µM cell strainers, and TILs were obtained by collecting the cells in the interphase after Ficoll (MP Biomedicals, #091692254) gradient separation. Spleens and livers were collected in ice-cold HBSS containing 2% FBS. Single cell suspension from spleens were generated through disruption on a wire screen (infection studies) or nylon nano mesh (tumor studies); Red blood cells were lysed with 0.84% (w/v) NH_4_Cl or ACK lysis buffer (Thermo Scientific). Livers were minced and digested with Collagenase IV deoxyribonuclease I in Hank’s balanced salt solution for 30 mins at 37 °C. Digested tissue was separated after centrifugation over Ficoll (MP Biomedicals, #091692254) and washed with complete media. Cells were resuspended in RPMI 1640 medium supplemented with 50 μM β-mercaptoethanol, penicillin (100 U/ml), streptomycin (100 mg/ml), and either 1 or 10% fetal bovine serum (FBS) or Click’s media (Irvine Scientific, #9195-500mL) containing 10% FBS, 10% Pen/Strep, and 10% Glutamine for flow cytometric analyses, as described below.

### Flow Cytometric Analyses

Flow cytometry data were collected on a BD FACSymphony with BD FACSDiva software or Invitrogen AttuneNXT cytometer with Attune cytometric software. All data were analyzed with FlowJo. Live-Dead staining was done using the aqua fluorescent reactive dye (ThermoFisher Scientific, L34966A) prior to surface staining. For surface and tetramer staining, cells were stained for 30-40 mins at 4°C. For transcription factor staining, cells were fixed and permeabilized with the eBioscience Foxp3/Transcription Factor staining kit (ThermoFisher Scientific, 00-5523-00) before staining with antibodies for 30-45 mins at 4°C. Antibodies used for flow cytometry included: anti-mouse Ly108 (330-AJ, Biolegend, 134608, 1:200), anti-mouse Tim3 (RMT3-23, Biolegend, 1:200), anti-mouse/human CD44 (IM7, Biolegend, 103026, 1:200), anti-mouse CD8a (53-6.7. BD, 563332, 1:200), anti-mouse CD8a (53-6.7, Invitrogen, 45-0081-82, 1:200), mouse anti-TCF7/TCF1 (S33-966, BD, 567018 1:100), anti-mouse CX3CR1 (SA011F11, Biolegend, 149048 & 149006, 1:200), anti-PD1 (RMP1-30, BD, 748266, 1:150), anti-KLRG1 (2F1, Invitrogen, 46-5893-82, 1:200), anti-CXCR6 (SA051D1, Biolegend, 151121, 1:200), anti-mouse CD45.1 (A20, Biolegend, 110743, 1:200), anti-mouse CD45.2 (104, Biolegend, 109824, 1:200), anti-mouse CD4 (RM4-5, Biolegend, 100526, 1:200), anti-mouse/Rat Foxp3 (FJK-16s, Invitrogen, 48-5773-82, 1:100), anti-mouse TCRβ (H57-597, Biolegend, 109251, 1:200), anti-mouse LAG3 (C9B7W, Biolegend, 125219, 1:200). H2-DbGP33 (1:200) and H2-DbNP396 (1:125) tetramers were acquired from the National Institutes of Health (NIH) Tetramer core at Emory University.

### Bulk RNA-seq

Freshly prepared splenocytes from day 30 LCMV CL-13 infected mice were sorted for bulk populations of GP33^+^ subsets using the BD FACSAria SORP Titan. Three to five mice were pooled for each replicate. Cells were homogenized in TRIzol LS reagent (Invitrogen, 10296028) and total RNA was isolated per protocol. Library preparation, quality control and paired end sequencing was performed at GeneWiz Inc RNA-seq data analysis was done as we previously described^(*79*)^. In brief, sequences were adapter and quality trimmed with Trim Galore (v0.6.10)^(*81*)^ and aligned to the mm10 reference genome^(*82*)^ using Star (v2.7.11a)^(*83*)^. Counts were generated with HTSeq (v2.0.5)^(*84*)^. Differential analysis was performed using edgeR (v3.42.4)^(*85*)^ and heatmaps were generated using limma (v 3.56.2)^(*86*)^.

### Assay for transposase-accessible chromatin with sequencing (ATAC-Seq)

ATAC-seq libraries for bulk sorted populations, as described above, were generated using previously published protocol^(*87*)^. Library quality control and paired end PE150 sequencing was performed by Novogene Co. For ATAC-seq analysis, reads were adapter and quality trimmed using Trim Galore (v 0.6.10)^(*81*)^. Trimmed fastq files were aligned to the mm10 reference genome^(*82*)^ with Bowtie2 (v2.4.3)^(*88*)^ using the following settings bowtie2 -p 10 --very-sensitive -k 10 -X 2000. Genrich (v0.6.1)^(*89*)^ was used to define peaks from biological replicates using the following settings –j –r -s 10 -v. Regions aligned to the mitochondrial chromosome and the Encode defined blacklist regions^(*90*)^ were excluded during peak calling with Genrich. Bedgraphs from Genrich were used to generate bigwig files using the UCSC utility bedGraphToBigWig^(*91*)^. Narrowpeak files for all samples were merged using BEDTools (v2.28)^(*92*)^, converted to the SAF format and counts were generated using featureCounts (v.2.0.6)^(*93*)^. Differential analysis was performed using edgeR (v3.42.4)^(*85*)^ and heatmaps were generated using limma (v 3.56.2)^(*86*)^. Genomic range objects were generated using GenomicRanges (v1.52.1)^(*94*)^ and annotated with ChipSeeker (v1.36.0)^(*95*)^. Pathway enrichment analysis was performed using ClusterProfiler (v4.8.3)^(*96*)^. Venn diagrams were generated with ggVennDiagram (v.15.0)^(*97*)^. HOMER (v4.11)^(*54*)^ was used for motif analysis. Genome tracks were viewed and generated with Integrative Genomics Viewer (v2.16.1)^(*98*)^.

### Statistical analysis

For all animal experiments, at least 3 mice were included in each group for every independent experiment. All experiments were repeated 2-5 times. Results are expressed as mean ± SEM. Data were analyzed using a two-sided Student’s t-test, one-way ANOVA, or two-way ANOVA after confirming their normal distribution. All analyses were performed using Prism 10.2.0 (GraphPad Software, Inc.) and p < 0.05 was considered statistically significant.

### Data Availability

The bulk RNA-seq and ATAC-seq data generated in this study will be deposited in the Gene Expression Omnibus (GEO) database. All deposited data will be publicly available, upon acceptance of the manuscript for publication. The remaining data in this study are provided with the manuscript, with source data provided herein.

## Acknowledgements

O.A.O. designed and performed experiments with mice and cells, analyzed data and wrote the manuscript. H.X. and J.T.I. did experiments with mice and cells. J.A.B, R.S.W, and A.J.Z. contributed to manuscript construction, data analyses, discussion, and manuscript editing. A.J.Z. provided viral stocks and working space. L.G. provided the Gfi1-tdTomato reporter mice. L.Z.S. was responsible for the original conceptualization of this study, overall data presentation, and manuscript construction; L.Z.S. acquired most of the funding for this study, designed experiments, supervised laboratory studies and data analyses, and edited the manuscript. All authors have met the requirements for authorship and are in consensus of the content in this publication. We acknowledge Dr. Leighton Grimes (Cincinnati Children’s Hospital) for help in acquiring the Gfi1-tdTomato mice as well as Sajesan Aryal from Dr. Rui Lu Lab (University of Alabama, Birmingham) for the protocol on sample preparation for ATAC-Seq. We thank other members of the Shi lab and the research team in the Department of Radiation Oncology for their constructive input. We are grateful for the Startup fund from the Department of Radiation Oncology and the O’Neal Invests pre-R01 Grant from the UAB-O’Neal Comprehensive Cancer Center to L.Z.S. This study is also partially funded by National Institutes of Health grants (1R21CA230475-01A1, 1R21CA259721-01A1, and 1R01CA279849-01A1 to L.Z.S.), the V Foundation Scholar Award (V2018-023 to L.Z.S.), a DoD-Congressionally Directed Medical Research Programs grant (ME210108 to L.Z.S.), a Cancer Research Institute CLIP Grant (CRI4342) to L.Z.S, and 5R01AI156290 to A.J.Z.

## Ethics Declaration

The authors declare no conflicts of interest.

---

**Figure S1.**
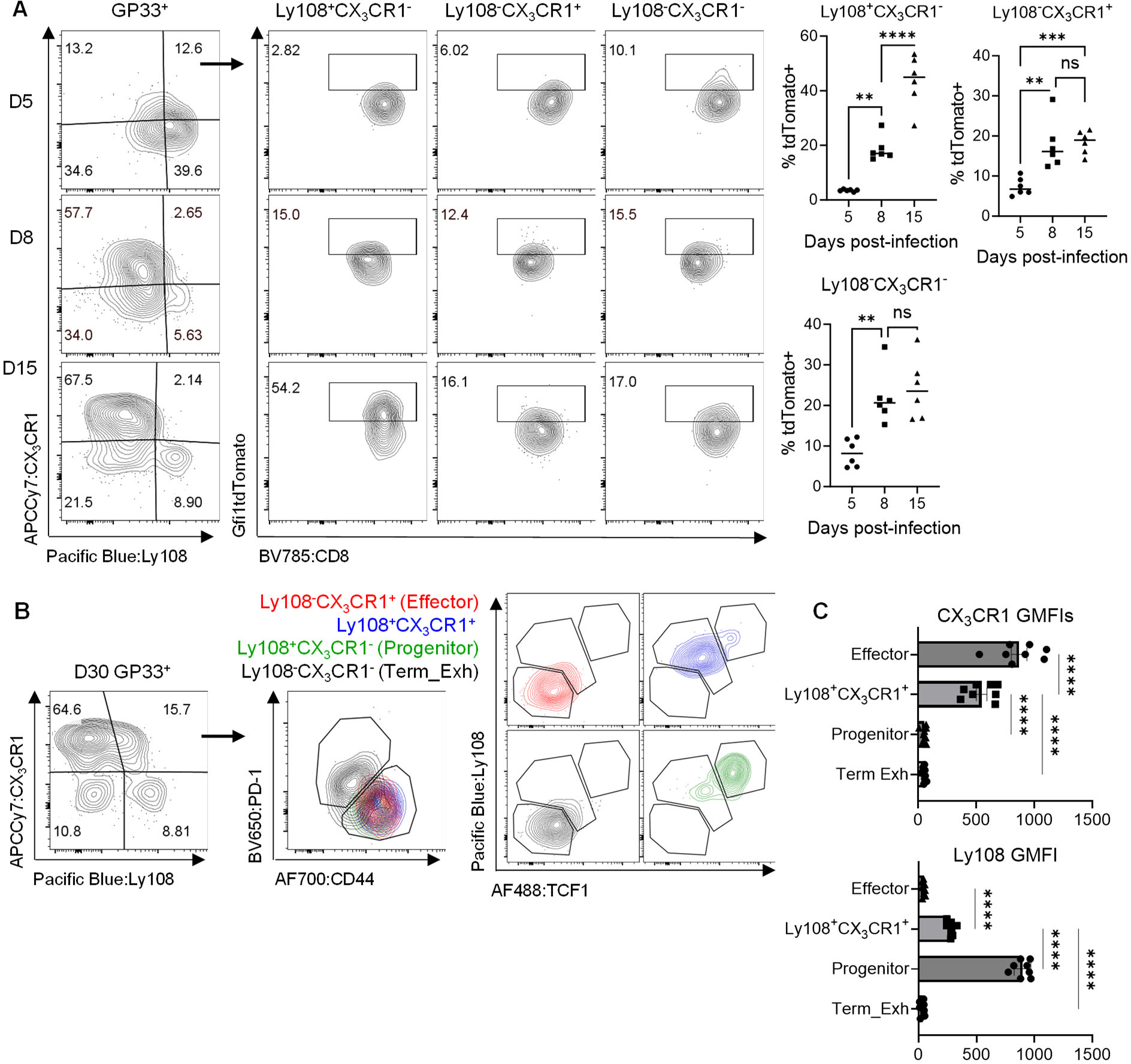
Gfi1 is differentially expressed in exhausted CD8 T cell subsets during chronic infection. **A.** Gfi1 (tdTomato) expression on GP33^+^ CD8^+^ T cells at different times post-LCMV Cl-13 infection. **B.** Expression of PD-1, CD44, Ly108, and TCF1 on the 4 subsets defined by CX_3_CR1 and Ly108. **C.** Flow cytometric analysis of CX_3_CR1 and Ly108 expression on LCMV specific subsets on day 30 post-LCMV Cl-13 infection. Data were from 2 independent experiments with n=7-8, shown as mean ± s.e.m, ns, P≥0.05; **, P<0.01; ***, P<0.001; ****, P<0.0001, analyzed by one-way ANOVA with Tukey’s or Sidak’s post-hoc test.

**Figure S2.**
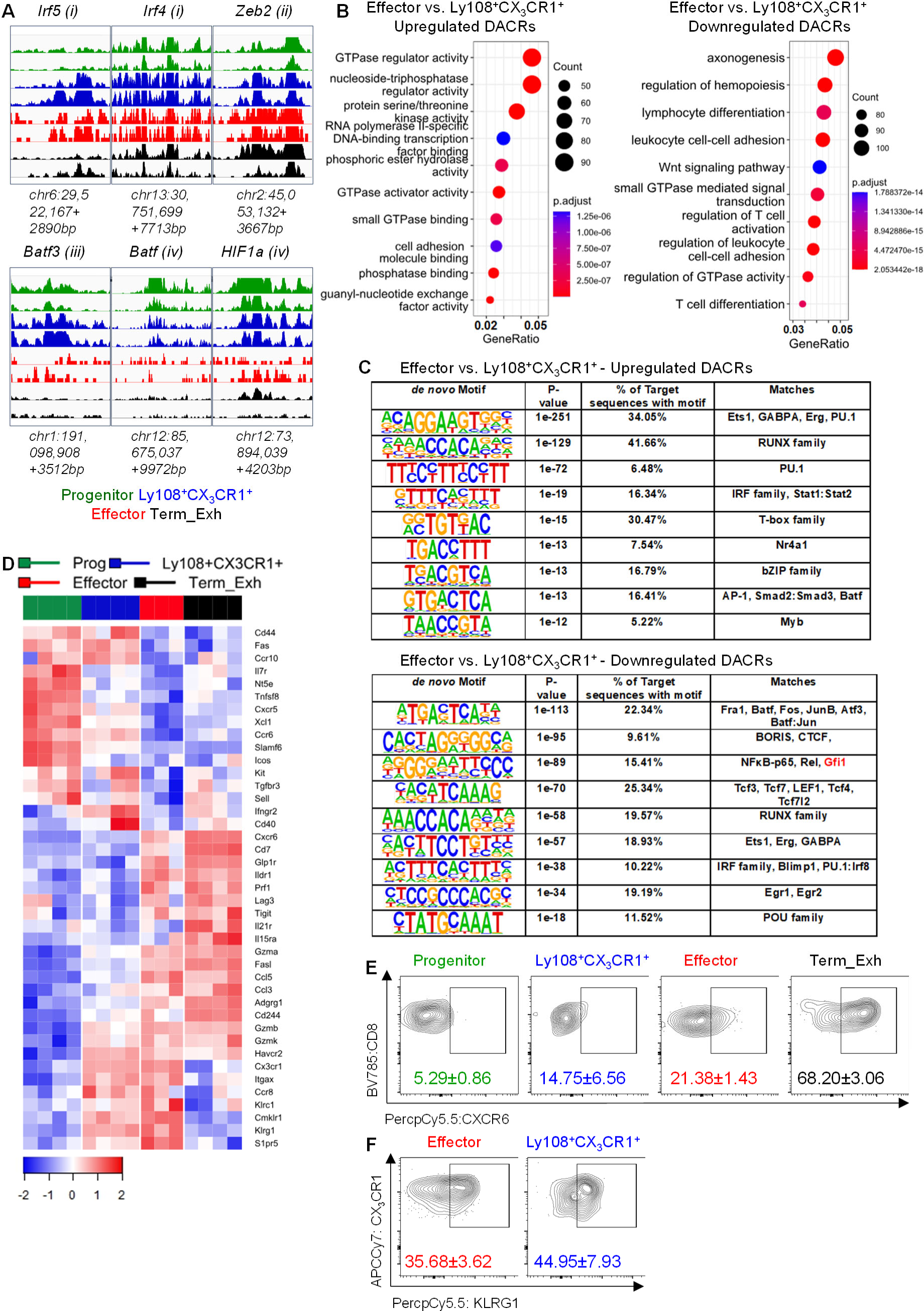
Ly108^+^CX CR1^+^ cells are a distinct subset with shared chromatin and transcription properties of progenitor and effector-like cells. GP33^+^ CD8^+^ T cells were harvested and sorted from LCMV Cl-13 infected WT mice on day 30 based on Ly108 and CX_3_CR1 expression for ATAC-Seq (**A-C**), and RNA-Seq (**D**). **A.** Genomic track view of accessibility profiles at select loci of the four subsets. **B.** GO enrichment pathway (Molecular Function) analysis of upregulated (left) and downregulated (right) DACRs in Ly108^+^CX_3_CR1^+^ and effector cells over progenitors. **C.** HOMER motif analysis of DACRs. **D.** Heatmap visualization of transcripts for select effector and activation molecules. Prog=Progenitor and Term_Exh=Terminally Exhausted. **E.** Flow cytometric quantification of CXCR6 expression on different subsets. **F.** Flow cytometric quantification of KLRG1 expression on Ly108^+^CX_3_CR1^+^ and effector cells. Flow data were representative of 2 independent experiments. ATAC and RNA-seq data were from 2 and 3-4 independent replicates, respectively. Each replicate was a pooled sample of 3-5 mice.

**Figure S3.**
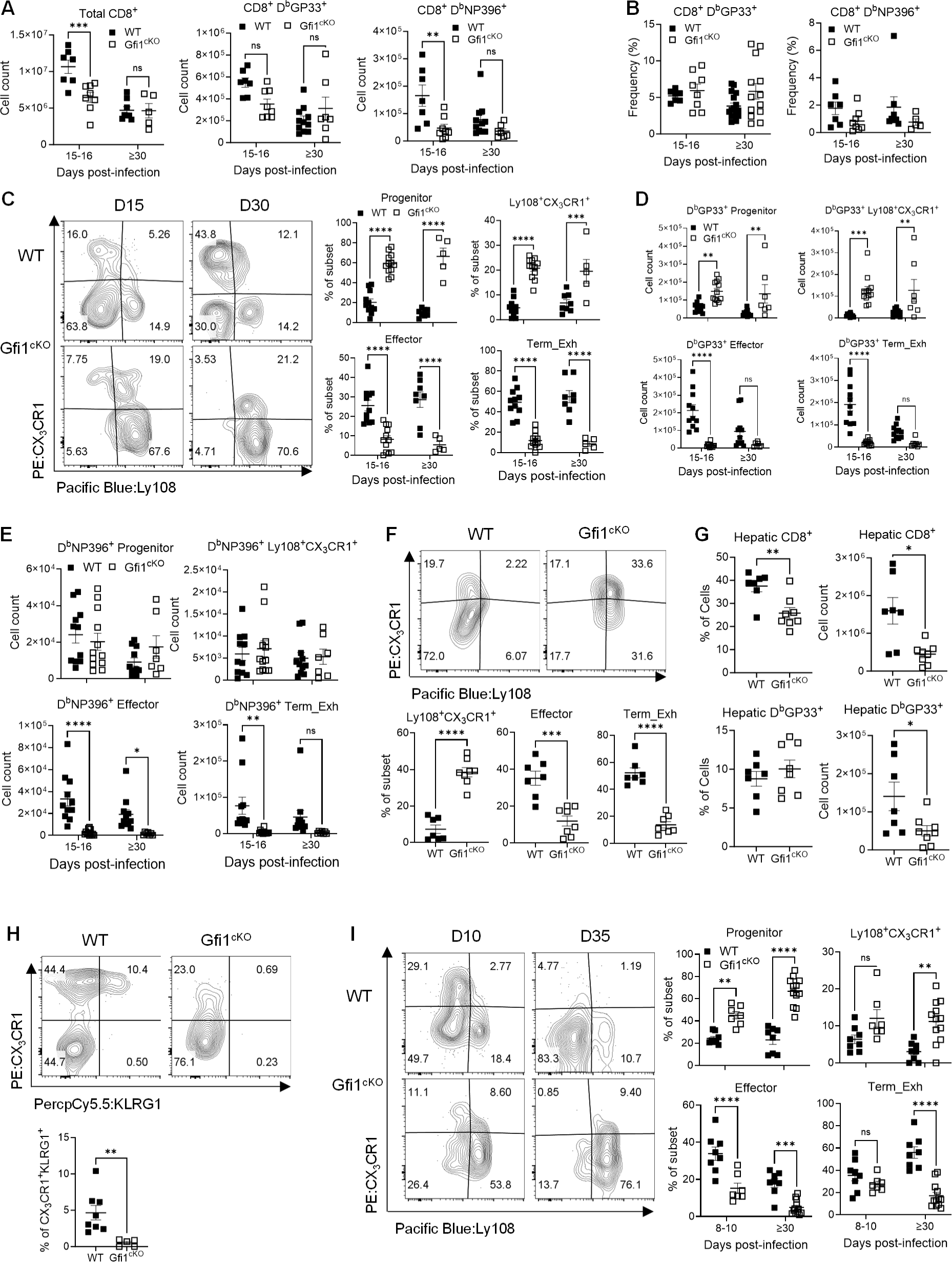
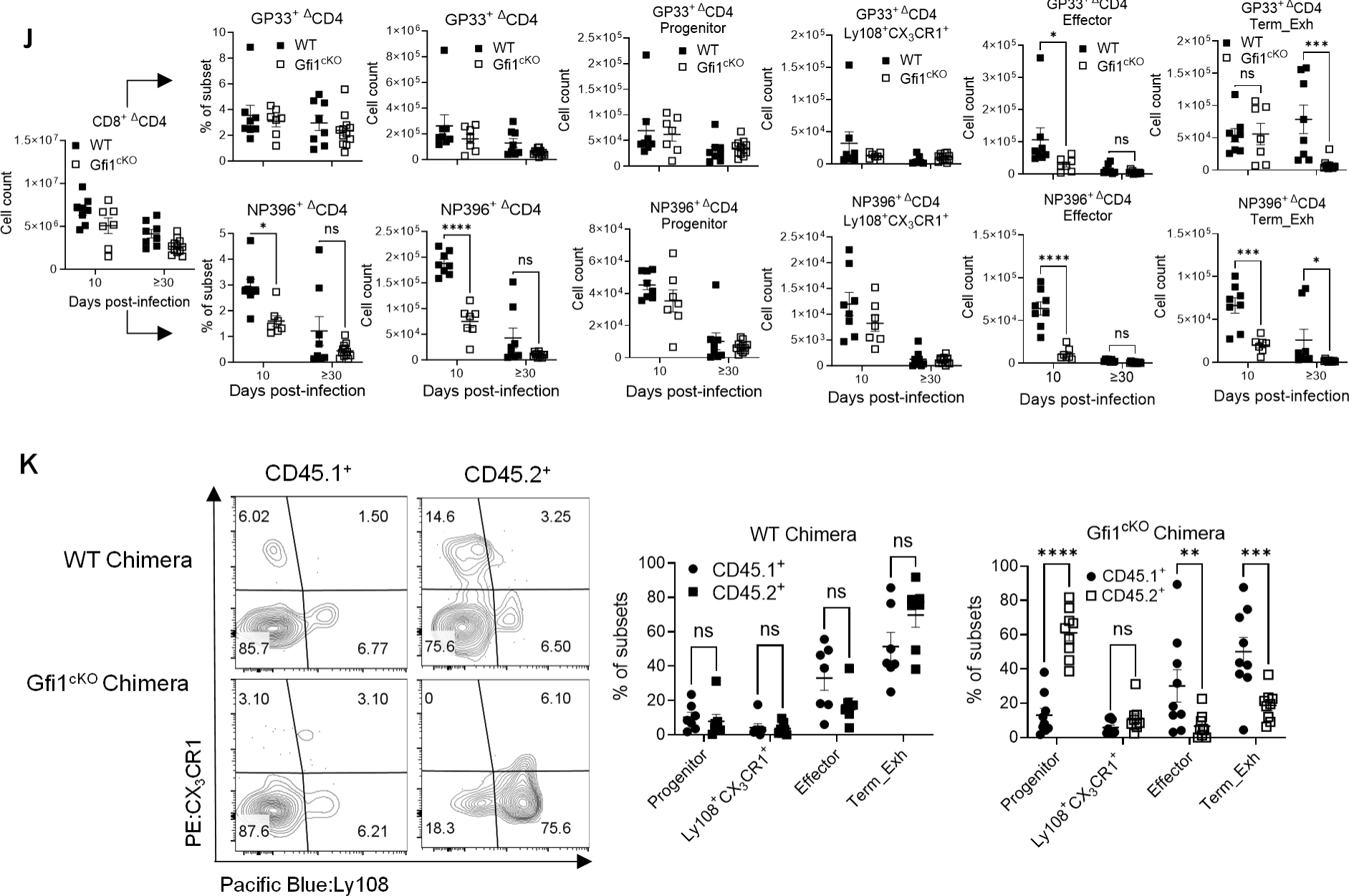
CD8^+^ T cell intrinsic Gfi1 controls the formation of effector and terminally exhausted subsets. **A-E.** WT and Gfi1^cKO^ mice were infected with LCMV Cl-13. At the designated time points, spleens were harvested for enumeration of total CD8^+^, GP33^+^ and NP396^+^ cells (**A**), frequencies of GP33^+^ and NP396^+^ cells (**B**), frequencies of the four subsets in NP396^+^ CD8^+^ T cells based on CX_3_CR1 and Ly108 expression (**C),** cell counts for all 4 subsets in GP33^+^ (**D**) and NP396^+^ CD8^+^ T cells (**E**). **F-G**. Livers from the LCMV Cl-13-infected mice were harvested on day 30 to evaluate CX_3_CR1 and Ly108 expression on GP33^+^ CD8^+^ cells (**F**), as well as the frequency and cell counts of total and GP33^+^ CD8^+^ T cells (**G**). **H.** KLRG1 expression in splenic NP396^+^ CD8^+^ T cells harvested on day 30 post-LCMV Cl-13 infection. **I.** Ly108 and CX_3_CR1 expression on NP396^+^ CD8^+^ T cells on day 30 post-LCMV Cl-13 infection with CD4^+^ T cells depleted. **J.** Cell counts of total CD8^+^ T cells, the frequency and cell counts of GP33^+^ and NP396^+^ CD8^+^ T cells, as well as cell counts of all 4 subsets in GP33^+^ and NP396^+^ CD8^+^ T cells from LCMV-infected WT and Gfi1^cKO^ mice depleted of CD4^+^ T cells. **K.** Ly108 and CX_3_CR1 expression on NP396^+^ CD8^+^ T cells from LCMV Cl-13 infected chimeras on day 30. Data were from 3 independent experiments, shown as mean ± s.e.m., with each dot denoting an individual mouse. ns, P≥0.05; *, P<0.05; **, P<0.01; ***, P<0.001; ****, P<0.0001. Data in **A-E** and **I-K** were analyzed with two-way ANOVA with Sidak’s post-hoc test. Data in **F-H** were analyzed with unpaired two-tailed Student’s t test.

**Figure S4.**
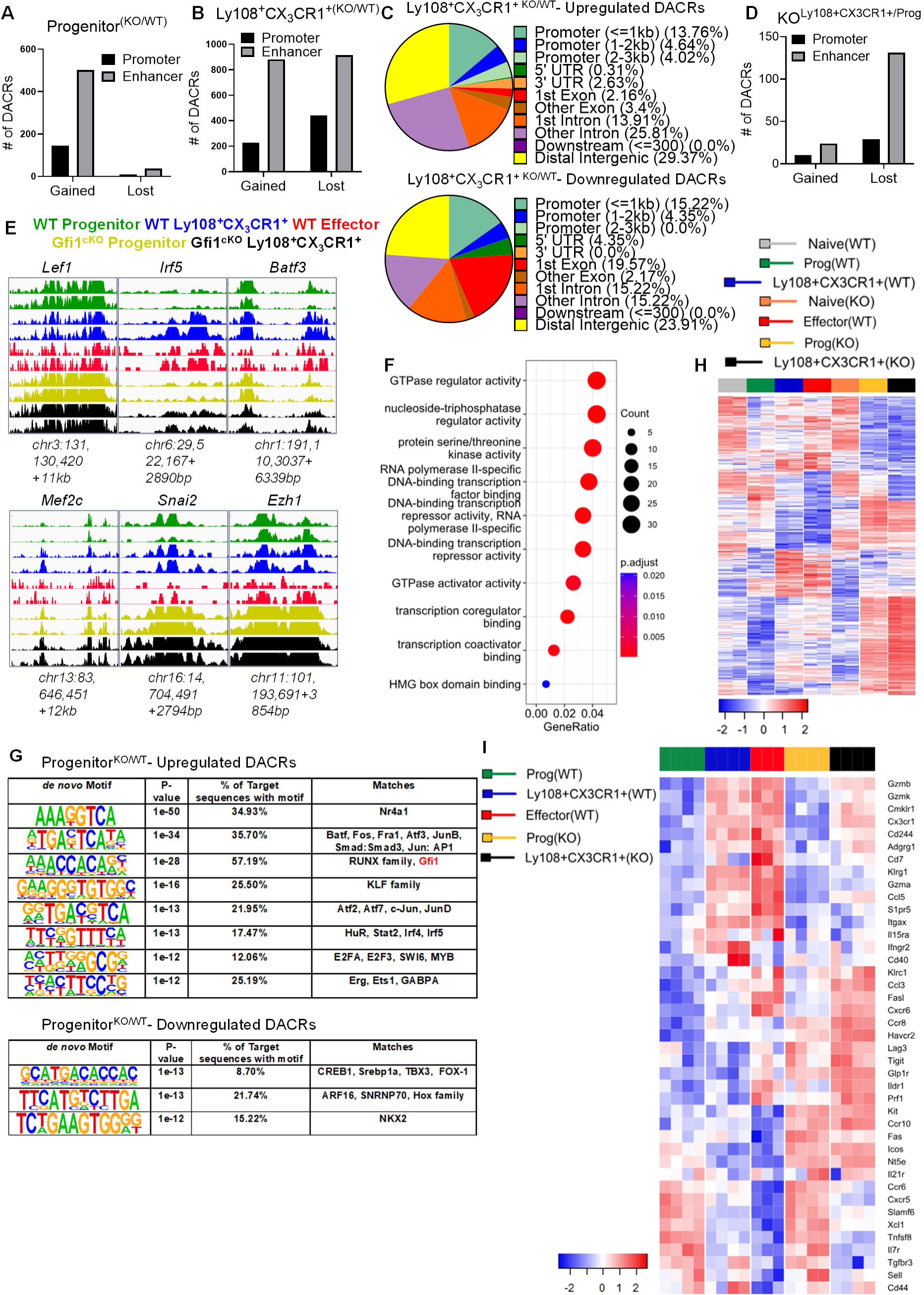
Gfi1 maintains the chromatin accessibility required for the formation of effector cells. GP33^+^ exhausted CD8^+^ T subsets were harvested and sorted from LCMV Cl-13 infected WT and Gfi1^cKO^ mice on day 30 for ATAC-seq (**A-H**) and RNA-seq (**I**). **A.** DACRs in Gfi1^cKO^ versus WT progenitors. **B.** Gfi1^cKO^ versus WT Ly108^+^CX_3_CR1^+^ cells. **C.** Distribution of DACRs between Gfi1^cKO^ and WT Ly108^+^CX_3_CR1^+^ cells. **D.** Gfi1^cKO^ Ly108^+^CX_3_CR1^+^ cells versus Gfi1^cKO^ progenitors. **E.** Genome track view of select DACRs in all the subsets. **F.** GO enrichment pathway (Molecular Function) analysis of upregulated DACRs in Gfi1^cKO^ versus WT Ly108^+^CX_3_CR1^+^ cells. **G.** HOMER motif analysis of DACRs. **H.** Heatmap visualization of DACRs between Gfi1^cKO^ and WT subsets with naïve cells included as controls. **I.** Heatmap visualization of transcripts for select TFs in WT and Gfi1^cKO^ GP33^+^ subsets. ATAC-seq and RNA-seq data were from 2 and 3-4 independent replicates, respectively. Each replicate was a pooled sample from 3 to 5 mice.

**Figure S5.**
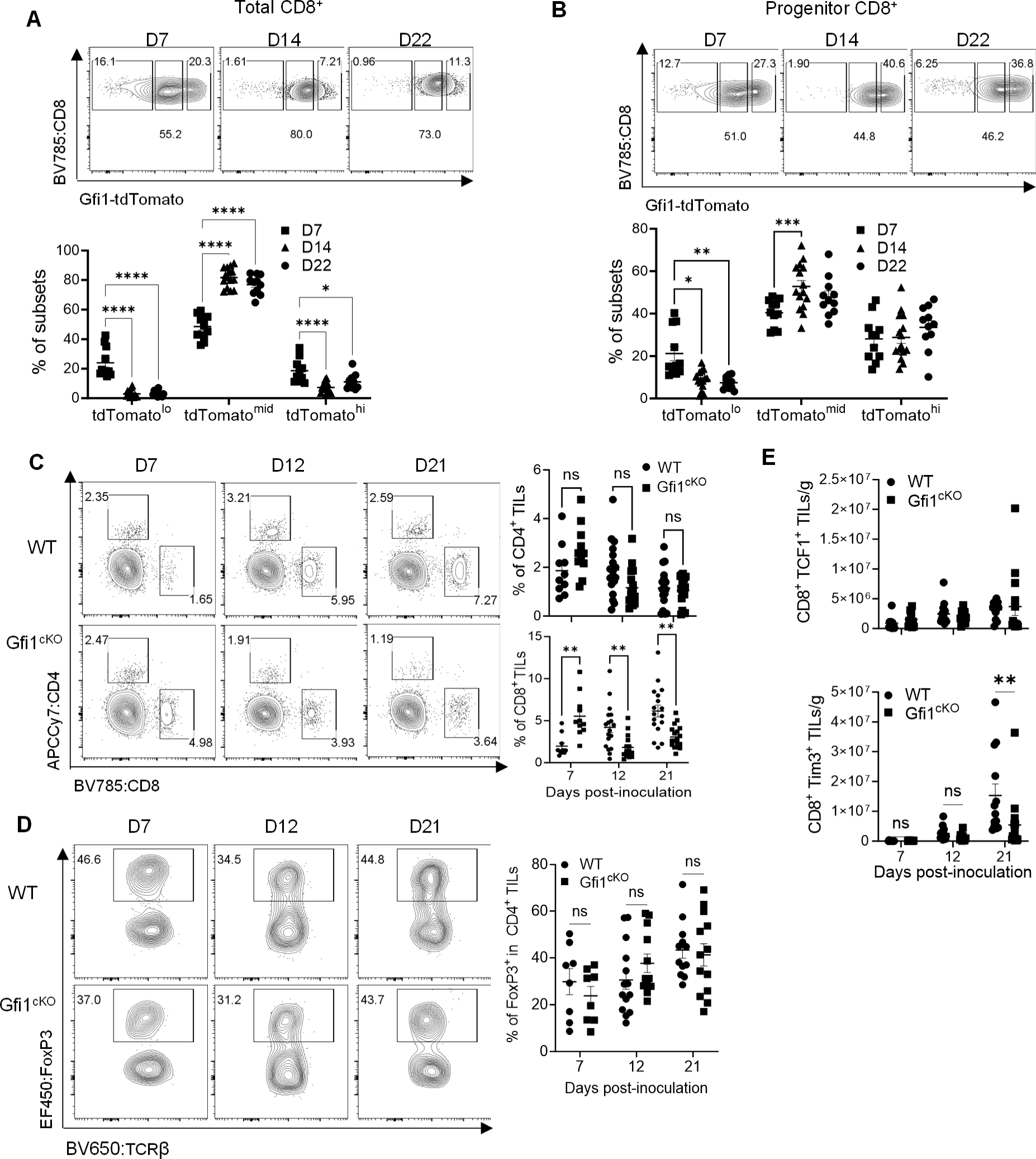
Gfi1 is required for the generation of terminally differentiated intratumoral CD8^+^ T cells. **A-B.** Gfi1^tdTomato^ reporter mice were inoculated with MB49 cells. At the designated times post-tumor inoculation, total CD8^+^ TILs (**A**) and TCF1^+^CD8^+^ TILs (**B**) were categorized to tdTomato^lo^, tdTomato^mid^, and tdTomato^hi^ cells, with their frequencies shown in the scatter dot plots (**A**). **C-E.** WT and Gfi1^cKO^ mice bearing MB49 tumors were euthanized at the designated times to determine the infiltration of CD4^+^ and CD8^+^ T cells (**C**), frequencies of T_reg_ among CD4^+^ TILs (**D**), and cell counts of TCF1^+^ (top panel) and Tim3^+^ CD8^+^ TILs (bottom panel) (**E**). Data were pooled results from two independent experiments, depicted as mean ± s.e.m., with each dot denoting an individual mouse. ns, P≥0.05; *, P<0.05; **, P<0.01; ***, P<0.001; ****, P<0.0001. Data were analyzed by two-way ANOVA with Tukey’s, Dunnett’s, or Sidak’s post hoc test.

**Figure S6.**
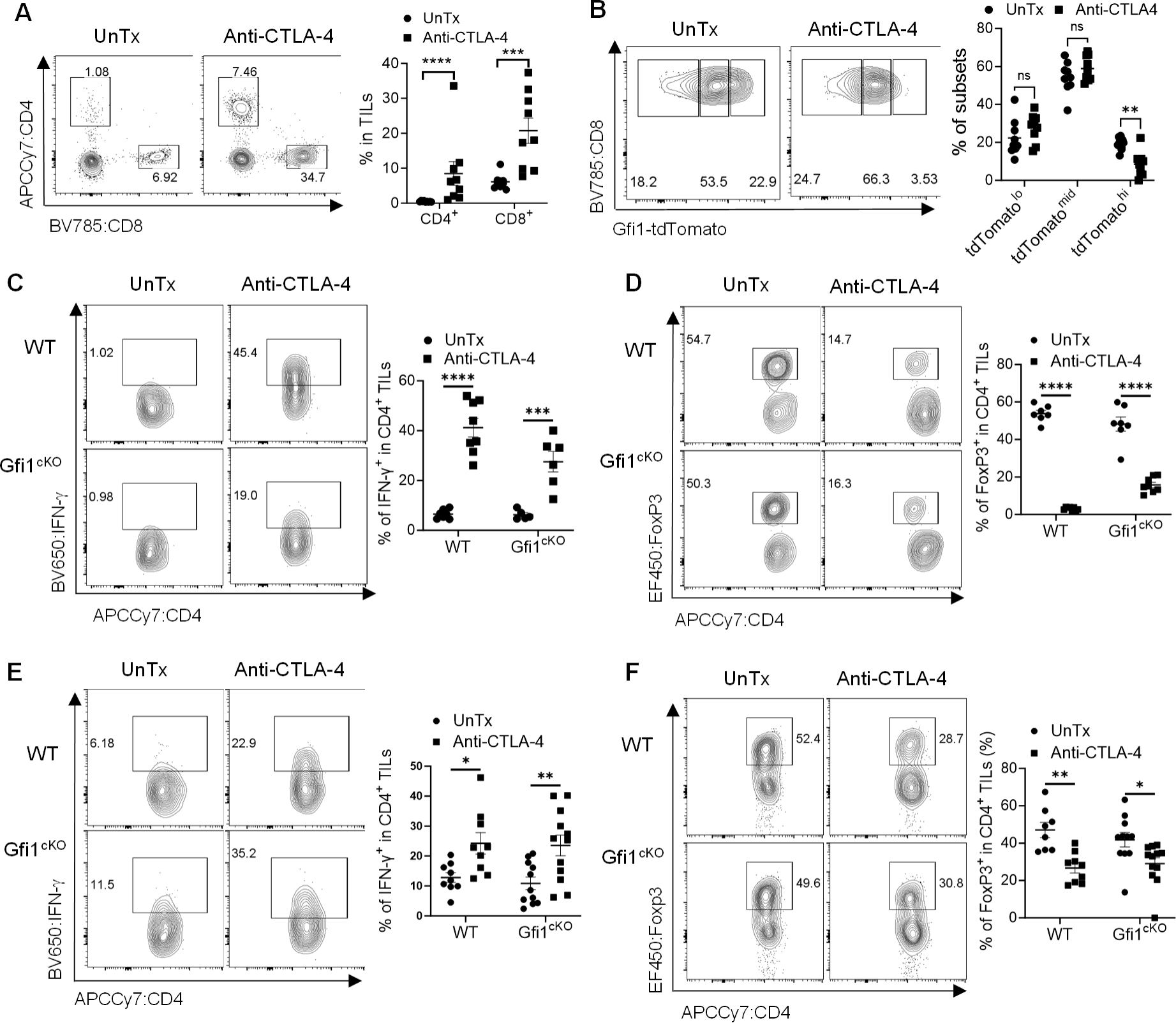
Gfi1 expression in T cells is largely dispensable for increased IFN-γ production and depletion of T_reg_ in CD4^+^ TILs by anti-CTLA-4. **A-B.** Gfi1^tdTomato^ reporter mice bearing palpable MB49 tumor were treated with or without anti-CTLA-4. Isolated single cell suspensions were analyzed for the frequencies of CD4^+^ and CD8^+^ TILs (**A**) as well as Gfi1-tdTomato expression in TCF1^+^CD8^+^ TILs (**B**). WT and Gfi1^cKO^ mice bearing palpable MB49 bladder tumor (**C-D)** or MC38 colorectal tumor (**E-F**) were treated with or without anti-CTLA-4. Isolated CD4^+^ TILs were analyzed for IFN-γ production (**C,E**) and the frequency of T_reg_ (**D,F**). Data were pooled results from two independent experiments, shown as mean ± s.e.m., with each dot denoting an individual mouse. ns, P≥0.05; *, P<0.05; **, P<0.01; ***, P<0.001; ****, P<0.0001. Data were analyzed by two-way ANOVA with Sidak’s post hoc test.

**Supplemental Figure 7.**
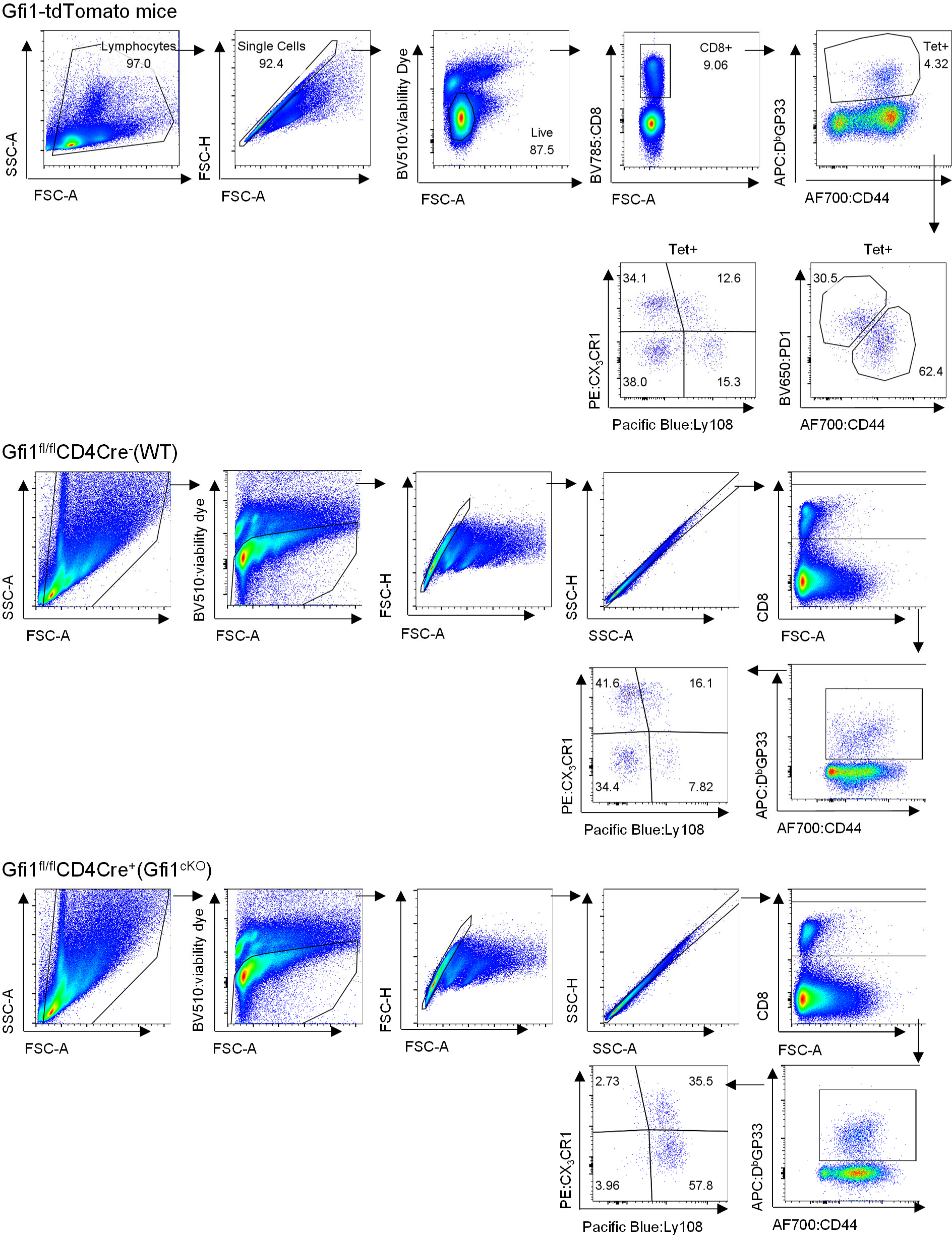
Main Gating Strategies.

